# Performance-impacting brain state maladaptation driving disease progression in early mouse and human neuroinflammation

**DOI:** 10.64898/2026.06.19.733346

**Authors:** Ting Fu, Kira Engeroff, Anna-Lena Schlegelmilch, Erik Ellwardt, Wei Fan, Michael Lippert, Tony Fernando de Schultz, Mona K. Roesler, Konstantin Radyushkin, Miriam Schillner, Manuela Ecker, Nicolas Ruffini, Anna Wierczeiko, Tim Hahn, Luisa Klotz, Michael J. Schmeisser, Frank W. Ohl, Frauke Zipp, Stefan Bittner, Albrecht Stroh

**Affiliations:** Department of Neurology, Focus Program Translational Neuroscience (FTN), Rhine Main Neuroscience Network (rmn2), University Medical Center of the Johannes Gutenberg University Mainz, Mainz, Germany; Department of Neurology, University Hospital Muenster (UKM), Muenster, Germany; Institute of Neuropathology, University Hospital Muenster (UKM), Muenster, Germany; Leibniz-Institute for Neurobiology, Magdeburg, Germany; Institute of Anatomy, Focus Program Translational Neuroscience (FTN), Rhine Main Neuroscience Network (rmn2), University Medical Center of the Johannes Gutenberg University Mainz, Mainz, Germany; Mouse Behavior Unit, Johannes Gutenberg University Mainz, Germany & Focus Program Translational Neuroscience (FTN), University Medical Center of the Johannes Gutenberg University Mainz, Mainz, Germany; Department for Psychiatry, Psychotherapy and Psychosomatic Medicine, University Medicine Halle, Martin-Luther University Halle-Wittenberg, Halle, Germany; Institute for Human Genetics, University Medical Center of the Johannes Gutenberg University Mainz, Germany; Institute of Translational Psychiatry, University Hospital Muenster (UKM), Muenster, Germany; Institute of Physiology I, University Hospital Muenster (UKM), Muenster, Germany

## Abstract

The neuronal mechanisms driving progression in neuroinflammatory disorders from early relapse-remitting phases to later neurodegenerative phases remain largely elusive. Functional brain state shifts towards hyperactivity, persisting beyond relapses, represent an early maladaptive response. Here, in remission stage of an experimental autoimmune encephalitis (EAE) mouse model of RRMS, we identified a reduced excitability upon optogenetic stimulation in the brain stem, the area of active disease, while in the cortex a persistent cortical neuronal hyperactivity and synaptic remodeling emerged, accompanied with an increase of markers of early apoptosis. In contrast, hippocampal circuits, which undergo a functional state shift without hyperactivity, do not show increased apoptosis. Visual cortical networks showed a deterioration of the accuracy of encoding visual information and a decrease in the behavioural visual discrimination ability in mice. In RRMS patients in remission, we identified a reduced visual colour discrimination, indicating both the presence and the clinical relevance of early brain state maladaptation that may contribute to progression independent from relapse activity (PIRA).

**Summary:** In a RRMS model and in patients, impaired visual processing was reported, indicating brain state maladaptations, associated with persistent cortical hyperactivity, brain stem hypoactivity, synaptic remodeling, and apoptosis. These maladaptations might contribute to relapse-independent disease progression through sustained network dysfunction.

## INTRODUCTION

Current therapeutic approaches in chronic CNS inflammation such as multiple sclerosis (MS) target inflammatory processes in the peripheral immune system and are effective at reducing clinical relapses and preventing new focal inflammatory-driven lesions (Wiendl et al., 2021) . However, most patients still suffer from diffuse neurological symptoms such as cognitive deficits, fatigue or anxiety (Murphy et al., 2017, Ellwardt et al., 2022, Di Filippo et al., 2018). These deficits seem to be rather independent of focal lesions and are associated with a worse long-term outcome. Indeed, over time, a substantial proportion of patients acquires a progressive worsening of neurological deficits, independent from clinical relapses, which is currently termed progression independent from relapse activity (PIRA). Recently, it has been shown that this process can start very early in the disease course in some individuals (Tur et al., 2023), and importantly, this process has been shown to drive the majority of disability accumulation in MS patients. Although inflammatory and neurodegenerative processes in MS are closely linked, the underlying pathological pathways determining i) the pathophysiology of PIRA in relapse-free phases of early-stage MS patients and ii) the later conversion into a predominantly progressive disease course are both poorly understood. In mouse models of early stages of neurological disorders, including the relapsing-remitting experimental autoimmune encephalomyelitis (EAE) model, early shifts in functional neuronal network architecture - brain state - have been observed (Ellwardt et al., 2018). These shifts, in different neurological disorders, occurred in cortical regions which were not directly affected by the underlying molecular or cellular pathophysiological events, including but not limited to the visual cortex (Arnoux et al., 2018, Ellwardt et al., 2018, Burgold et al., 2019, Jafari et al., 2021a, Zott et al., 2018, Rosales Jubal et al., 2021, Luttjohann et al., 2026). Notably, while these early maladaptations are mainly characterized by the emergence of hyperactive neurons, an early hypoactive phenotype has also been reported in a focal cortical EAE model (Jafari et al., 2021a). Early brain state shifts might be brought about by plasticity mechanisms, as a homeostatic reaction to an anatomically distant molecular or cellular event, aiming for retaining global network stability (Stroh et al., 2024). Yet the interdependency between the emergence of hyperactive neurons in primarily non-affected brain regions with subsequent neurodegeneration and its phenotypic representation remains elusive. Indeed, MS patients report subtle neurocognitive symptoms as well as fatigue also at early disease phases (Disanto et al., 2018) or even in the prodromal phase of the disease (Menascu et al., 2019). In a synergistic and longitudinal experimental design, in mouse and man, we here reveal the transitions of individual neurons to hypo- and hyperactive states. We determine the consequences of neuronal hypo- and hyperactivity on local cortical and hippocampal network function. To elucidate a chain of events functionally linking the initial neuroimmunological event, i.e. neuron - T-cell interaction to distant primarily unaffected cortical networks, we investigated the excitability of brainstem neurons, a region which is prominently affected in EAE. We assess early signs of cell death as consequence of hyperactivity, and excitatory and inhibitory synapse densities. Lastly, to evaluate the translational relevance of early brain state maladaptations in remission stage, we tested discrimination and object recognition in 20 stable and mildly affected Relapsing Remitting Multiple Sclerosis (RRMS) patients. These findings should inform network-rebalancing interventions with the prospect of re-balancing maladaptive and persistent brain states and preventing activity-dependent neurodegeneration and progression independent from relapse activity.

## RESULTS

### In EAE relapses, both hypo- and hyperactive neurons emerge, followed by a hyperactive-only network state in remission

We employed longitudinal awake two photon (2-P) calcium imaging of head-fixed mice on a jet ball, allowing for the assessment of locomotion while imaging cortical microcircuit activity through a cranial window (Grienberger and Konnerth, 2012). For mimicking the human RRMS, we subjected the mice to an EAE paradigm, focusing on the first peak of the disease. Prior to the onset of the first peak, mice displayed normal running behaviour, without sign of discomfort or stress (Fig. 1A, Suppl. Fig. 1). For a fine-grained assessment of the dynamics of local microcircuit activity with single-cell resolution, we conducted longitudinal 2-P calcium imaging in the awake behaving mouse. We restricted the expression of the genetically encoded calcium indicator GCaMP6f to excitatory neurons only, upon identifying excitatory neurons as the source of plasticity-mediating TNFα in our previous study (Ellwardt et al., 2018). We focused our microcircuit assessments on layers II / III of the primary visual cortex (V1). We injected CamKII-GCaMP6f AAV into the visual cortex and implanted a chronic window (Fig. 1A) about four weeks prior to the induction of EAE. This enabled to longitudinally image mice at three given time points: 1. before EAE induction (= pre-induction); 2. at EAE peak score (relapse); 3. in remission, defined as the absence of significant motoric disease symptoms. In the *in vivo* awake behaving 2-P imaging, stable and strong expression of GCaMP6f in layer II / III of the mouse visual cortex was observed and individual cells could be identified at all three imaging time points (Fig. 1B-D). We employed somatic 2-P calcium imaging as a proxy of suprathreshold neuronal activity measures, i.e. the spiking output of the given neuron (Grienberger and Konnerth, 2012). First, we assessed spontaneous, ongoing, neuronal activity in awake behaving mice in V1. At pre-induction, the cortical microcircuit exhibited typical sparse ongoing activity of excitatory pyramidal neurons in layer II / III. The silent cell proportion increased in relapse and decreased in remission to levels similar to the pre-induction phase (Fig. 1E). In relapse, the animals exhibited significant motor symptoms, such as tail and hind limb paralysis. On the level of cortical microcircuit activity, we did not observe a significant increase in the mean activity rate in relapse when compared to pre-induction (Fig. 1F). In remission, the firing rate increased significantly (Fig. 1F). It is noteworthy, that while the mean firing rates in relapse were not significantly altered in comparison to pre-induction, the variance of the firing rates significantly increased throughout the course of the disease, from pre-induction, to relapse, to remission, indicating a stratification of firing rates (Fig. 1G). Indeed, when assessing the mean firing rates on the level of the individual animals, we observed a rather diverse effect in relapse, followed by a clear increase of mean firing rates across all animals in remission (Fig 1K). In contrast, in a longitudinal assessment of network states in healthy control mice, no significant alteration in network state occurred throughout the three time points (Suppl. Fig. 2A-D).

**Figure 1:**
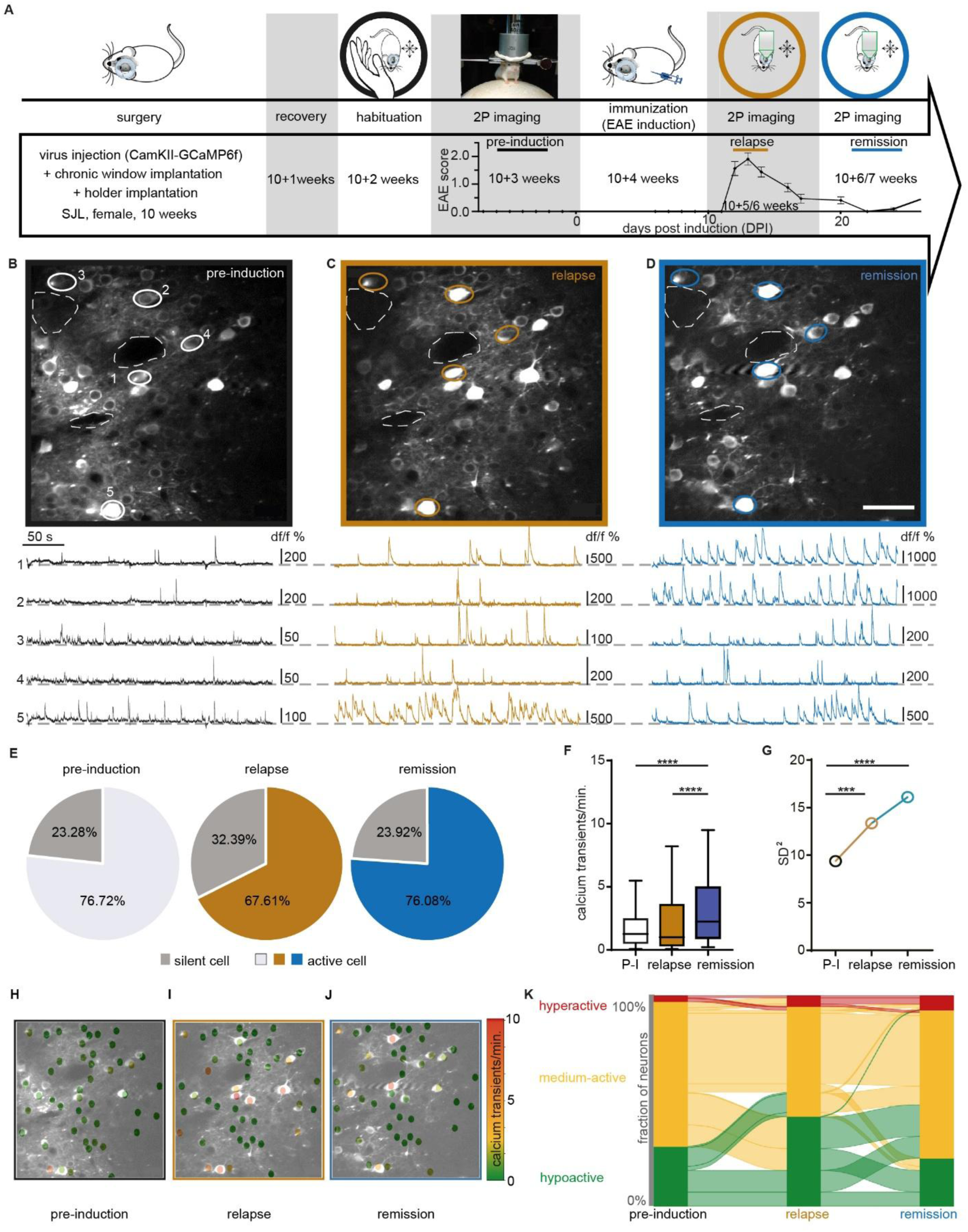
Temporal emergence of cortical network reorganization revealed by *in vivo* longitudinal 2-P calcium imaging of awake behaving mice. **A:** Experimental paradigm of *in vivo* longitudinal 2-P calcium imaging in an EAE mouse model and corresponding EAE disease course with indication of imaging time points. **B - D**: 2-P imaging (upper) and corresponding calcium transients (below) of identical neurons in layer II + III of primary visual cortex of an awake behaving mouse at three different time points of disease: **B** pre-induction (black), **C** relapse (brown), **D** remission (blue). Scale bar =100µm. **E:** Silent cell proportion from all recorded cells at three different time points of disease; **F**: Average calcium transients per minute in P-I, RL and RM. Median: P-I=1.27, RL=1.00, RM=2.24; *p* _P-Iv.s.RL_>0.05, \*\*\*\**p* _P-Iv.s.RM_<0.0001, \*\**p* _RLv.s.RM_ =0.0088, one-way ANOVA test. **G**: Variance of the rate of calcium transients. Significant increase of variance in RL and RM in comparison to P-I. SD^2^_P-I_=9.38, SD^2^_RL_=13.36, SD^2^_RM_=16.10, ***P_P-Iv.s.RL_=0.0005, ****P_P-Iv.s.Rm_=0.0005, P_RLv.s.Rm_=0.2595 (Levene’s test for the equality of variance). **H - J:** Overlay of 2-P micrograph with a depiction of the neuronś activity level (indicated by the color scheme). **K:** Sankey/Alluvial plot showing the activity transition of all cells in all time points (hyperactive/red, medium-active/yellow, hypoactive/green). *n* = 5 animals and 267 neurons.

We next subdivided neurons across all animals based on their activity levels (Fig. 1H-J). We categorized the cells, according to their firing rate, into hypoactive (lowest 25%), medium-active and hyperactive (highest 2%) at pre-induction phase (Ellwardt et al., 2018). While on average, the activity rates in relapse were not yet altered, we now unravel an increase of the fraction of hypoactive neurons in relapse, paralleled by a modest increase in the fraction of hyperactive neurons (Fig. 1K). In remission, the fraction of hypoactive cells returned to pre-induction levels, while the increased fraction of hyperactive cells increased further (Fig. 1K). However, the overall significant shift of the mean activity rates in remission was not due to a further increase of hyperactive cells, but, to a transitioning of previous hypoactive cells (in relapse) back to medium-active cells (Fig. 1K).

### Synchronicity and connectivity of excitatory cortical networks break down in remission

Functional microcircuit imaging does not only allow the longitudinal assessment of individual neurons over time, but also reveals the signature of local network activity, i.e. the level of coordination or synchronization of the activity of individual neurons to each other, in a bound network (Melloni et al., 2007, Singer, 2013). At pre-induction, we identified the typical compartmentalized synchronicity motives, with clusters of increased synchronicity within the local microcircuit (Fig. 2A, D). In relapse, the local synchronicity clusters decreased sharply. This breakdown in local connectivity worsened further in remission, where the local synchronicity remained at a significant lower level compared to pre-induction (Fig. 2A - G, Suppl. Fig. 3A). We now specifically interrogated the impact of activity state on the connectivity matrix. We asked, which activity class contributes most to local network connectivity. Surprisingly, we found, that the medium-active cells carry the main fraction of the coordinated activity of the network (Fig. 2H, I, Suppl. Fig. 3B). Consequently, while the network increases the fraction of medium-active cells when transitioning from relapse to remission, this is not sufficient to counterbalance the detrimental effect of hyperactivity to local network signature. These findings again point to a maladaptive nature of the brain state shift, at local network scale, underlying efficient information processing (Aedo-Jury et al., 2020).

**Figure 2.**
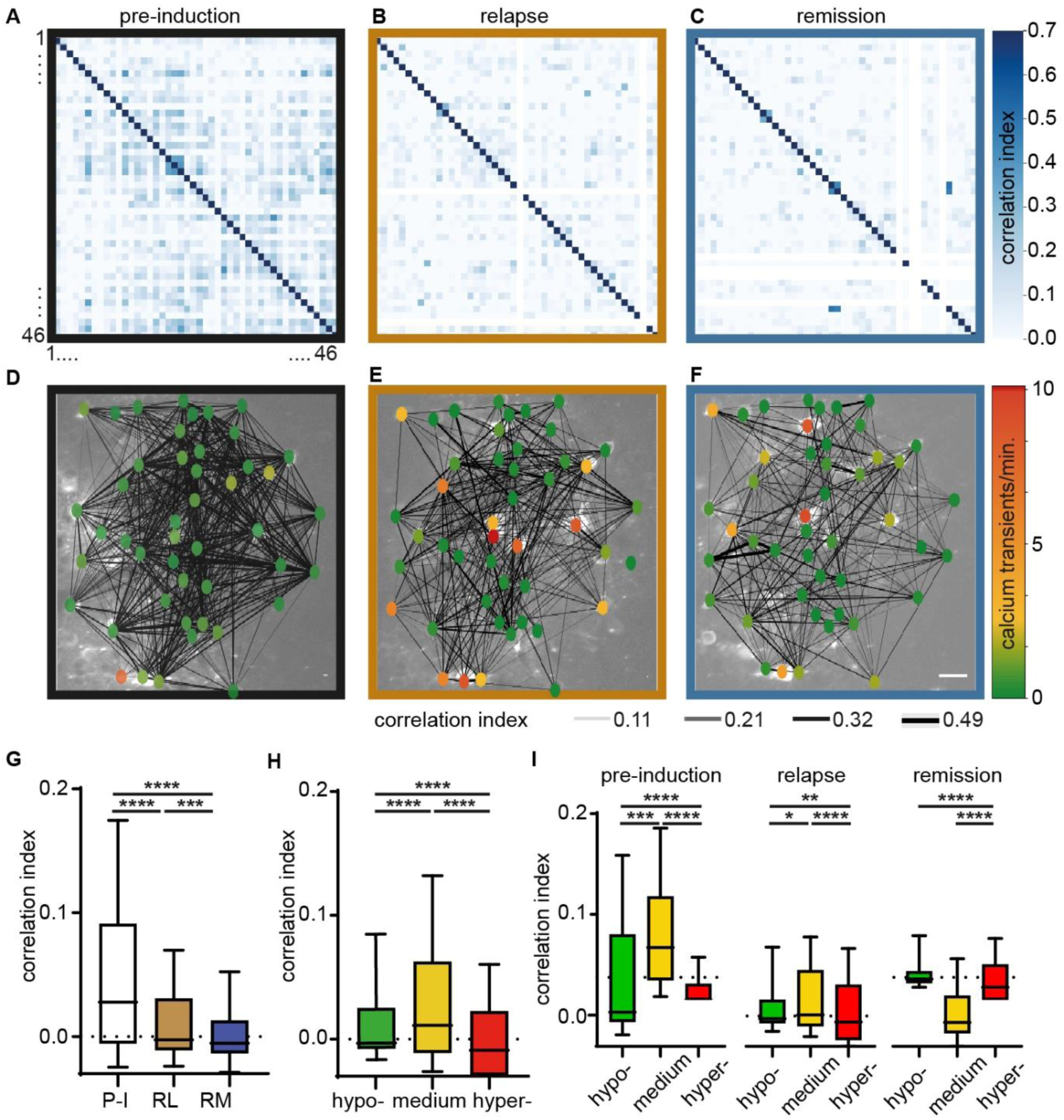
Synchronicity and connectivity of excitatory neuronal activity break down in relapse and remission phase of EAE. **A - C**: Synchronicity matrices of the identical local cortical microcircuit. **A** pre-induction (P-I), **B** relapse (RL), **C** remission (RM). **D - F**: Overlay of 2-P micrograph with a depiction of the neuron’s activity level (indicated by the color scheme) and connectivity (indicated by the width of the lines). **D** P-I, **E** RL, **F** RM. **G:** Correlation indices differ significantly between the time points. Median: P-I=0.028, RL=-0.0026, RM=-0.0053; ****p _P-Iv.s.RL_<0.0001, ****p _P-Iv.s.RM_<0.0001, ***p _RLv.s.RM_=0.0007, one-way ANOVA test. **H:** Correlation indices also differ significantly between the activity levels of all time points combined. Median: hypo-active group=-0.0029, medium-active group=0.011, hyper-active group=-0.0090; ****p_hypo-v.s.medium-_<0.0001, p _hypo-v.s.hyper-_<0.0001, ****p_medium-v.s.hyper-_ <0.0001, one-way ANOVA test. **I:** Sub-grouped correlation indices differ significantly between the activity levels in different time points. Median: pre-induction: hypo-active group=-0.0039, medium-active group=0.035, hyper-active group=-0.034; ***p_hypo-v.s.medium-_ =0.0004, ****p _hypo-v.s.hyper-_<0.0001, ****p_medium-v.s.hyper-_<0.0001, relapse: hypo-active group=-0.0026, medium-active group=0.0012, hyper-active group=-0.0059; *p_hypo-v.s.medium-_=0.0420, **p _hypo-v.s.hyper-_=0.0014, ****p_medium-v.s.hyper-_<0.0001, remission: hypo-active group=-0.0041, medium-active group=-0.0063, hyper-active group=-0.0143; p_hypo-v.s.medium-_=0.9957, ****p _hypo-v.s.hyper-_<0.0001, ****p_medium-v.s.hyper-_<0.0001, one-way ANOVA test. *n* = 46 neurons.

### Network state shift devoid of hyperactivity in hippocampal circuits

To compare the changes observed in the neocortex to those in other brain regions, we recorded single unit activity from the hippocampus using silicon probes. Since the between-group design does not allow for longitudinal observations of cells across days, we instead compared the population spike rates across the duration of the recording and between groups. While average firing rates across the whole recording did not show a statistical difference (binomial test, p=0.34), the distributions of short-duration (1 s) activity states over time were clearly different between the EAE remission group and pre-induction controls (Fig. 3A). A quantitative QQ-plot analysis revealed that the significant difference between the two distribution is caused by a higher fraction of neurons in the EAE group occupying medium-rate quantiles (Kolmogorov-Smirnov test, p<<0.01, Fig. 3B). Thus, the statistical effect of significant reduction of occupation in both the low-rate and high-rate intervals is camouflaged by a mean-value analysis. This also becomes evident by plotting the firing-rate data similar to Fig. 1K, as a fraction-of-activity plot (Fig. 3C). These findings indicate that EAE mice exhibit altered network dynamics in the hippocampal formation, with a significantly increased fraction of medium-rate firing, in sharp contrast to the functional state shift in cortex.

**Figure 3.**
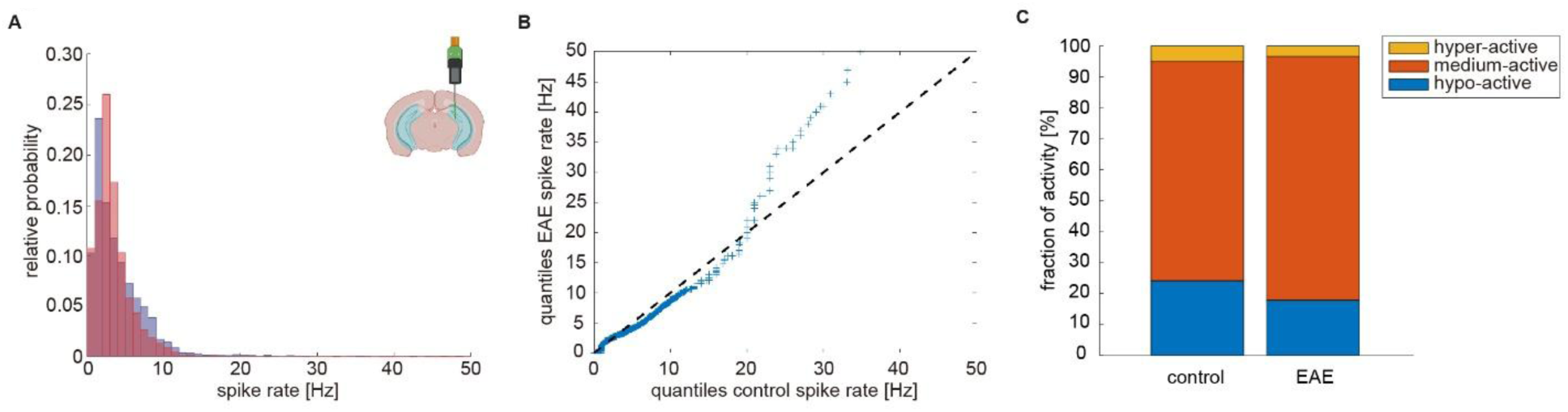
Distribution of spike rates in the hippocampus between pre-induction and EAE remission differ significantly. **A:** Distribution of spike rates for EAE (red) and the control group (blue). **B:** Quantile-quantile plot of spike rates of EAE group against pre-induction control (Kolmogorov-Smirnov test, p<<0.01). The dashed line indicates quantiles when EAE and control group quantiles would derive from the same distribution. **C:** Change in network structure characterized by an increase in the relative fraction of medium-active neurons.

### No apparent signs of inflammation in the cortex

Next, to exclude the possibility of a localized inflammation due to the injection of GCaMP6f significantly contributing to local network maladaptive reorganization, we histologically assessed inflammation at the injection area and compared this to the contralateral non-injected hemisphere. To also exclude systemic effects of EAE induction, we added a separate control group of non-immunized mice (Suppl. Fig. 4). We found only very few CD4 positive T cells, mostly adjacent to the meninges. Most importantly, we did not observe a CD4 infiltration in the area of GCaMP6f injection or in any other cortical areas. Along this line, we detected similar amounts of Iba-1 positive microglia in both hemispheres and almost no Mac-3 positive activated macrophages. Mice that underwent EAE showed no apparent signs of cortical immune cell infiltration or microglia activation neither in the area of injection nor in the contralateral hemisphere compared to healthy mice.

### Numbers of inhibitory synapses are increased in relapse and remission in layers II - IV / in visual cortex

We asked, whether the functional brain state shift identified by neurophysiological methods is reflected in neuroanatomical changes on synaptic level. We assessed the numbers of excitatory and inhibitory synapses in the visual cortex across all cortical layers (Fig. 4, Suppl. Fig. 6). We performed immunofluorescent staining for excitatory (VGlut1, Homer) and inhibitory (VGat, Gephyrin) synapses and quantified clusters (Fig. 4 A-D). Inhibitory synapses in the visual cortex were significantly increased in EAE relapse and remission compared to pre-induction animals (Fig. 4 E-F). However, only in layer II/III and IV this finding was significant, in layer V and VI a trend towards increased inhibitory clusters was detected (Fig. 4F). Strikingly, excitatory synapses did not change during the disease development. The assessed presynaptic and postsynaptic clusters can be appreciated in Fig. 4 and Suppl. Fig. 6. Interestingly, these changes of inhibitory synapses were found in a region with absent immune cell infiltration as stated already above, pointing again at an active network plasticity process.

**Figure 4.**
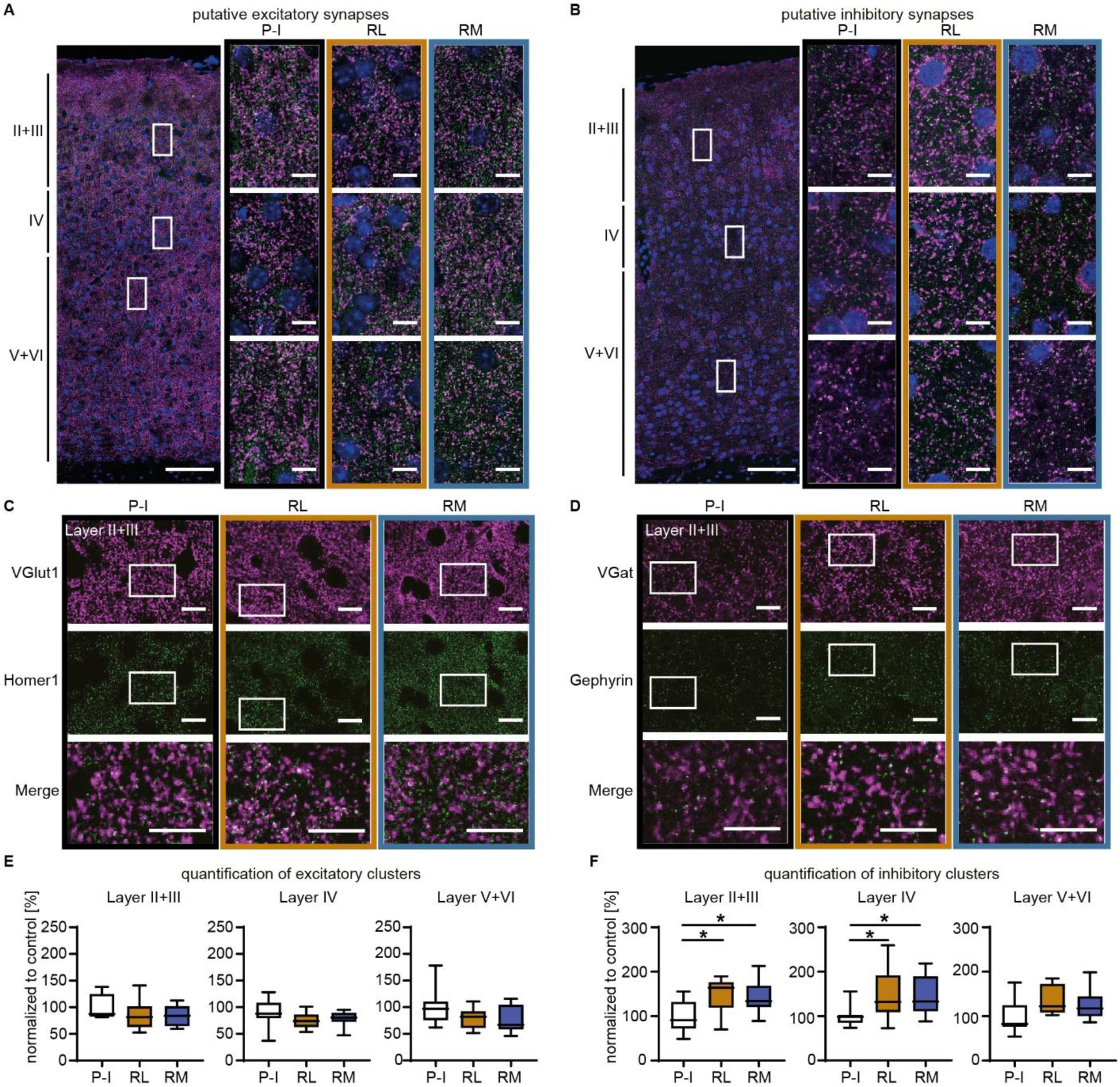
Number of inhibitory synapses are increased in layer II+III and layer IV in visual cortex area V1. **A**: Left (1^st^ column): overview of the V1 area in pre-induction (P-I) mice (green = Homer1, magenta = VGlut1, blue = DAPI). Rectangles indicate sites of example pictures shown on the right side. Right (2^nd^ – 4^th^ column): example pictures of Layer II+III, Layer IV and Layer V+VI of the V1 region in P-I (lack frame), relapse (RL) (brown frame) and remission (RM) mice (blue frame). **B**: Left (1^st^ column): overview of the V1 area in P-I mice (green = Gephyrin, magenta = VGat, blue = DAPI). Rectangles indicate sites of example pictures shown on the right side. Right (2^nd^ – 4^th^ column): example pictures of Layer II+III, Layer IV and Layer V+VI of the V1 region in P-I (lack frame), RL (brown frame) and RM mice (blue frame). **C**: Example pictures of pre- and postsynaptic clusters in P-I, RL and RM mice in Layer II / III of V1. The 3^rd^ row shows merged images which represent excitatory synapses (selections are indicated in the pictures above). **D**: Example pictures of pre-and postsynaptic clusters in P-I, RL and RM mice in Layer II / III of V1. The 3^rd^ row shows merged images which represent inhibitory synapses (selections are indicated in the pictures above). **E**: Quantification of excitatory synapses in Layer II + III, Layer IV and Layer V + VI shows that synapse numbers are not different compared to the synapse number of P-I mice of each Layer, respectively. **F**: Quantification of inhibitory synapses in Layer II + III, Layer IV and Layer V + VI shows that inhibitory synapses in the visual cortex were significantly increased in RL and RM mice compared to P-I animals, but only in layers II - IV (layer II + III mean ± SD: P-I 100.0 ± 36.2, RL 146.0 ± 39.1, RM 143.6 ± 35.8; pre-induction vs. relapse *p = 0.023 and pre-induction vs. remission 0.032; layer IV: pre-induction 100.0 ± 23.8, relapse 152.0 ± 57.2, remission 147.4 ± 43.9; pre-induction vs. relapse *p = 0.028 and pre-induction vs. remission 0.046; n = 3 animals and 9-10 slices for each condition, one-way ANOVA test)

### Decreased excitability in EAE relapse in excitatory brainstem neurons

To shed light on the mechanisms by which the brain state changes in primarily unaffected cortical and hippocampal circuits are linked to the disease-defining pathophysiology, i.e. neuron - T-cell interaction, we conducted all-optical-physiology experiments (Fu et al., 2021) in the brain stem, the main spot of inflammation in EAE.

The injection of rAAV1-Syn.GCaMP6f, to assess neuronal activity, and rAAV2-CamKII.C1V1.mcherry, for optogenetic stimulation of excitatory neurons, into the dorsal brainstem took place at least 3 weeks before calcium 2-photon imaging (Fig. 5A). Therefore, EAE induction was performed two weeks after viral transduction. To assess whether cells during EAE might not respond to optogenetic stimulation because they underwent apoptosis we performed caspase 3 staining and evaluated the percentage of GCaMP6 / caspase-3 double positive cells. Here we could not find a difference between groups (Fig. 5B, C). The co-expression of both GCaMP6 and C1V1 neurons was similar between both groups (Fig. 5D). The GCaMP6 density at the dorsal brainstem was not significant altered (Fig. 5E). EAE animals presented first disease symptoms after 7 days and reached score 1 from day 12 on (Fig. 5F).

**Figure 5:**
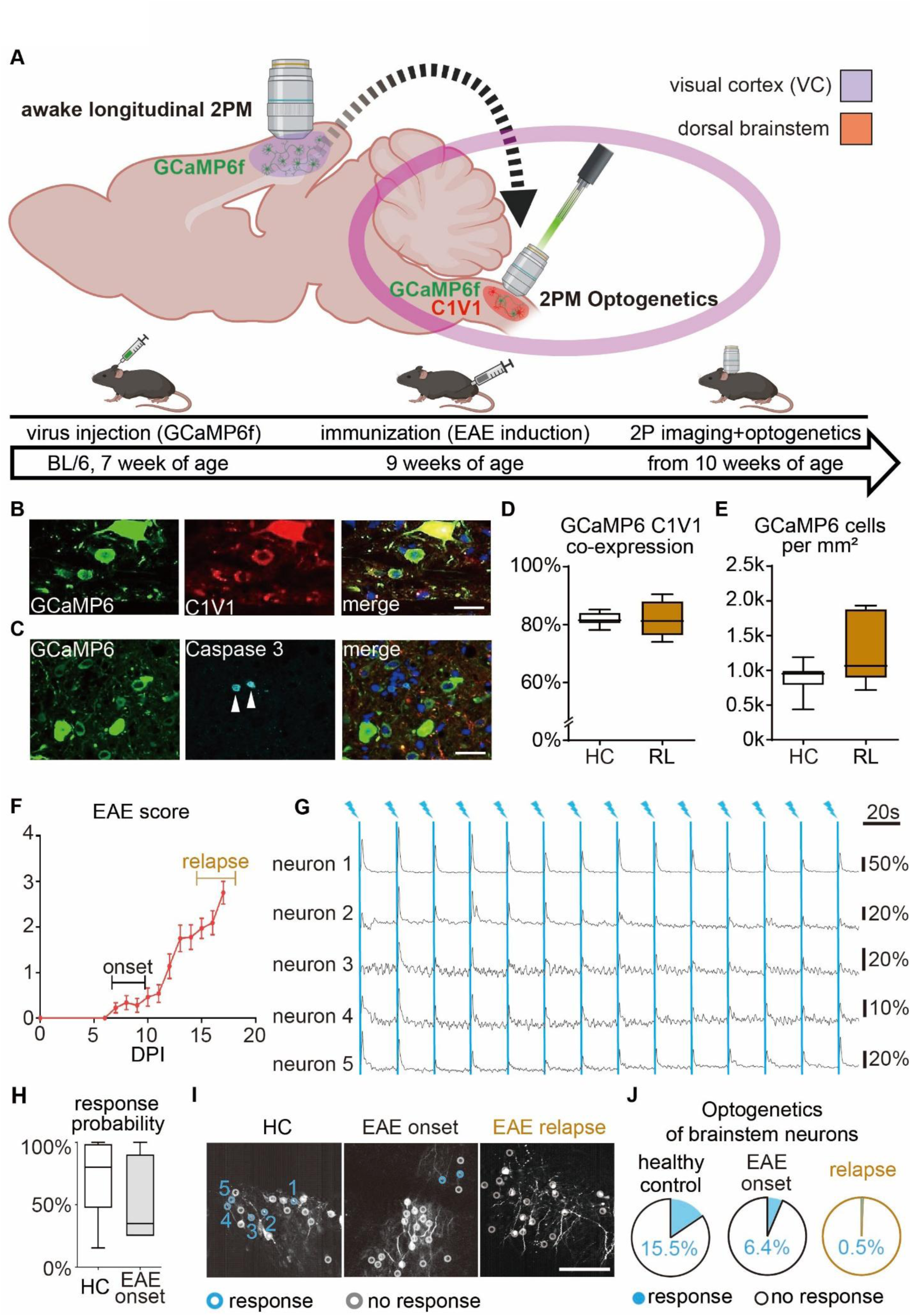
Reduced excitability upon optogenetic stimulation in the brainstem: **A:** Scheme of *in vivo* all optical physiology experimental timeline combining two-photon calcium imaging with optogenetics in EAE. The stereotactical injection of AAvs encoding GCaMP6 and C1V1-mCherry into the brainstem occurred at least 3 weeks before imaging and EAE induction was done two to three weeks after viral transduction. **B and C:** confocal imaging of GCaMP6, C1V1-mCherry, Caspase 3 and overlay including nuclear staining (DAPI). Scale bar = 25µm. **D and E:** No significant differences between control and EAE relapse for GCaMP6 / C1V1 co-localization (D, EAE_mean_: 0.824 ± 0.023, Control_mean_: 0.818 ± 0.009) and GCaMP6 density (E, EAE: 12672 ± 1802 cells per mm², Control: 8788 ± 891 cells per mm²). The co-expression of both proteins ranged above 80% (n = 3 animals and 7 slices each group, p > 0.05, unpaired two-sided students t-test). **F**: EAE time course, EAE pre-onset was defined with a score > 0 and < 1 and EAE relapse with a score ≥ 2, dpi = days post immunization. n = 20 animals **G:** Calcium transients recorded from the brainstem of a control mouse with optogenetic stimulation every 20s. All 3 displayed neurons show a calcium transient following each light pulse. **H:** The fraction of evoked transients following light stimulation was not different between control and EAE pre-onset (p = 0.209, Mann-Whitney test). Error bars represent standard error of the mean (SEM). **I:** Representative *in vivo* two photon micrograph with GCaMP6f transfected neurons (gray) and color-coding (blue) for responiveness following a light pulse (λ=552nm). Scale bar = 50µm. **J:** Fraction of neurons which respond at least to one light pulse during the recordings. Significant decrease of photoreactive neurons during EAE compared to controls (X² < 0.001 for controls vs. EAE relapse and X < 0.01 between EAE pre-onset and EAE relapse, controls (n = 3 animals and 129 neurons), EAE pre-onset (n = 3 animals and 109 cells), EAE peak (n = 9 animals and 196 neurons).

Optogenetic stimulation was applied to GCaMP6/C1V1 double transduced neurons (Fig. 5G). The response probability upon optogenetic stimulation did not differ significantly between EAE pre-onset and the healthy control group (Fig. 5H). Notably, the response rate dropped significantly during EAE pre-onset and almost vanished in EAE relapse were observed (Fig. 5I, J). These results indicate the presence of an initial brain state shift towards decreased excitability in those regions affected by inflammation.

### Sign of early cell death in areas of hyperactive neurons

While cortical neurons in EAE mice displayed a shift towards hyperactivity, in the hippocampus, we observed an increased fraction of neurons with medium activity in the network spike rate in EAE mice. We now asked, in a comparative experimental design, whether neuronal hyperactivity induced by EAE-mediated shift in neuronal state is linked to subsequent neurodegeneration. To identify neuronal cell death in EAE, we performed Fluoro Jade C (FJC) staining. To ensure the feasibility of the staining procedure, MCAO (Middle Cerebral Artery Occlusion) mouse tissue was assessed as positive control. Compared to MCAO tissue, the EAE tissue showed no positive signal of FJC staining. In remission, we did not identify an enhanced rate of neuronal cell death (Suppl. Fig. 7). Consequently, we needed to explore a marker involved in early signaling leading to subsequent cell death. Using RNAscope, we discovered changes in mRNA levels of the cell death regulator Receptor-interacting serine/threonine protein kinase 1 (RIPK1) (Fig. 6). RIPK1 mRNA is significantly increased in visual cortex of EAE mice (Fig. 6A-D) but not in hippocampus (Fig. 6E-L).

**Figure 6.**
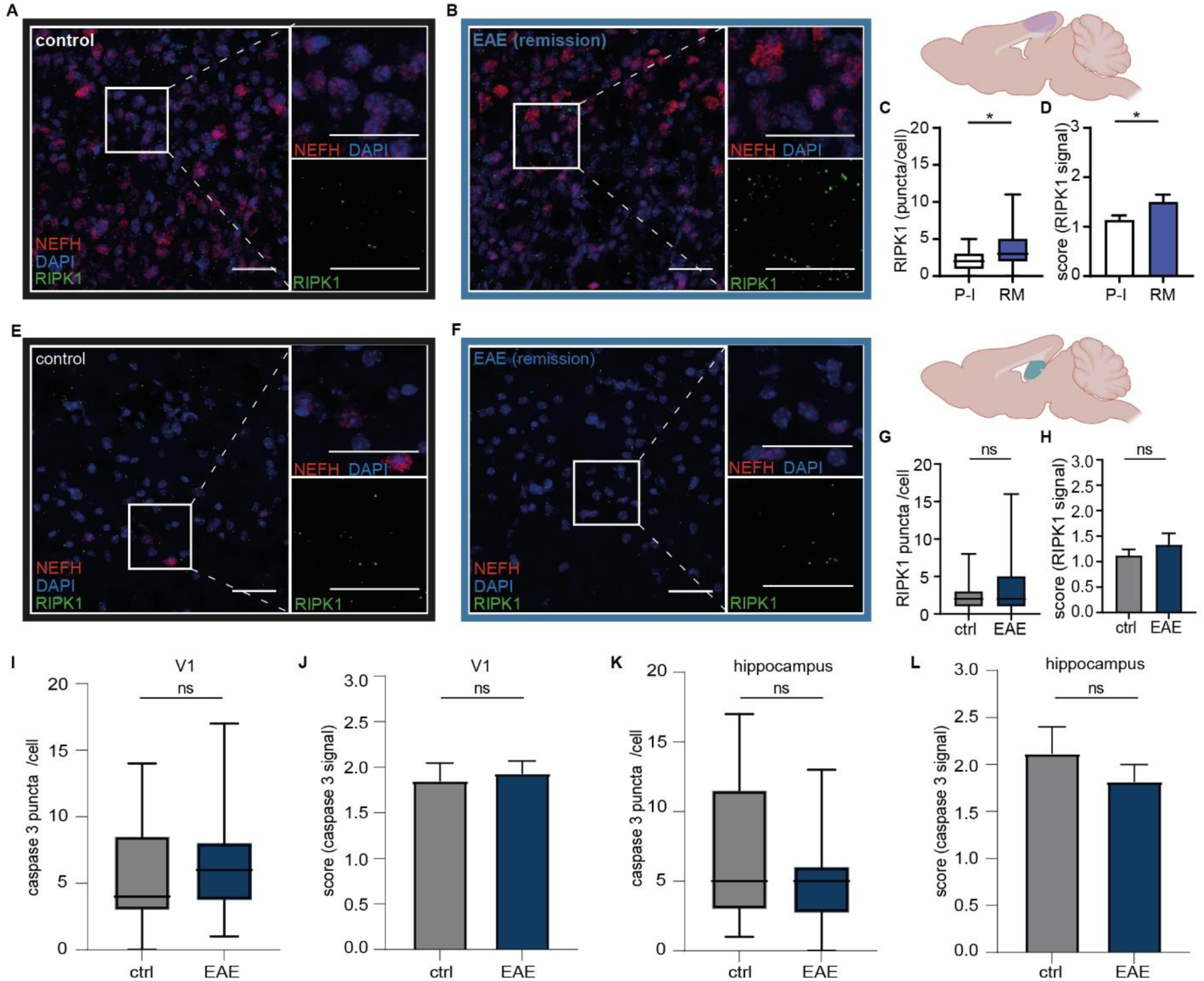
Early sign of cell death is shown in visual cortex but not hippocampus in EAE remission mice. **A, B:** Representative image of RNAscope for RIPK1 (green), NEFH (red) and DAPI (blue) in the visual cortex of control (A) and EAE (B) animals. **C, D:** Quantification of visual cortex RIPK1 signal in control (n=3) and EAE remission (n=3). **C.** RIPK1 puncta were counted in NEFH positive cells. (P= 0.0138; *P < 0.05). **D.** RIPK1 was quantified based on defined signal intensity scores (P= 0.0485; *P < 0.05). **E, F:** Representative image of RNAscope for RIPK1 (green), NEFH (red) and DAPI (blue) in the Hippocampus of control (E) and EAE (F) animals. **G, H:** Quantification of hippocampal RIPK1 signal in control (n=3) and EAE remission (n=3). **G**. RIPK1 puncta were counted in NEFH positive cells. (P= 0.5075). **H**. RIPK1 was quantified based on defined signal intensity scores (P= 0.4585). **I - L:** Quantification of Caspase 3 signal in control (n= 2) and EAE remission (n= 3). **I**: Caspase 3 puncta were counted in NEFH positive cells of the visual cortex. (P= 0.3199). **J**: Caspase 3 was quantified based on defined signal intensity scores in the visual cortex (P= 0.8076). **K**: Caspase 3 puncta were counted in NEFH positive cells of the Hippocampus (P= 0.3480). **L**: Caspase 3 was quantified based on defined signal intensity scores in the Hippocampus (P= 0.6967). Scale bars 50 µm. Box-and-whisker plot indicates the median value (centerline), the 25–75th percentiles (box) and the maximum and minimum (whiskers). P values were determined using two-tailed Mann-Whitney U test. Error bars represent standard error of the mean (SEM).

### Visual discrimination in EAE is significantly impaired in remission on both functional microarchitecture and behavior level

While indeed, spontaneous activity seems to be a very sensitive marker of the functional microarchitecture we now asked, whether the main function of primary visual cortex circuitry, i.e. the representation of visual afferents, is affected. For that, we subjected awake head-fixed mice to static and drifting gratings, provided by the 270° monitor ring, while recording local circuit function (Fig. 7A, B). Since the pioneering experiments of Hubel and Wiesel almost 60 years ago (Hubel and Wiesel, 1962), we know that excitatory neurons in layer II / III specifically and selectively react to distinct orientations and directions of drifting gratings. Prior to immunization, in pre-induction we observed a typical pattern of orientation and direction sensitive neurons, identified based on their selective response upon a specific stimulus (Fig. 7C - I). Particularly the polar plots highlight the selective responses (Fig. 7C - E). In relapse and remission, the visuotopic responses ceded drastically, as demonstrated by a broadening response pattern in the polar plot, and by a significant reduction of both orientation selectively and direction selectivity indices. Both disease stages did not differ in terms of their lack of the ability of local circuits to adequately represent these key features of the visual environment (Fig. 7F, G).

**Figure 7:**
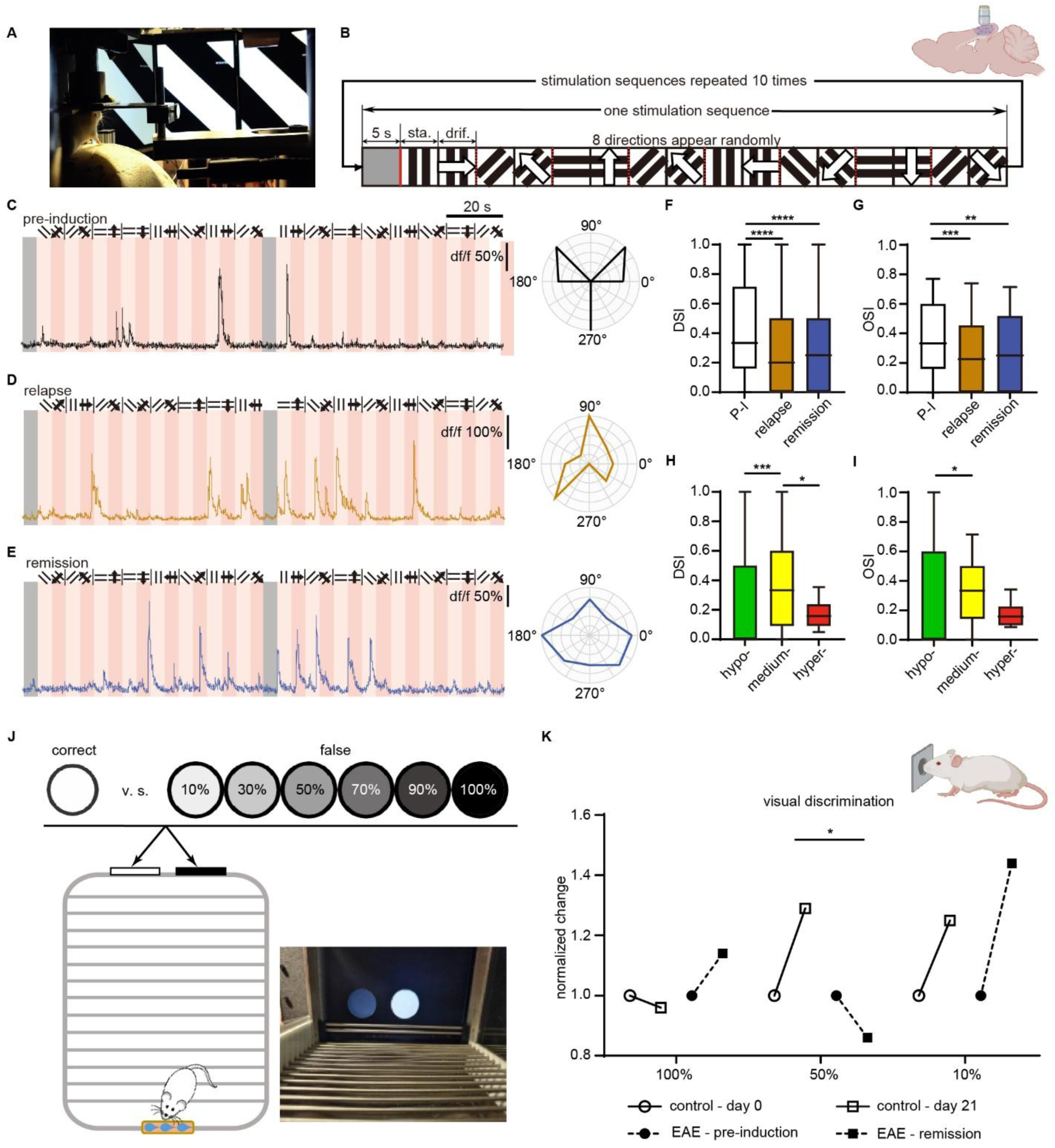
Visual discrimination in EAE is significantly impaired in remission in both functional microarchitecture and behaviour level. **A:** Image of an awake head-fixed mouse in the virtual reality system during 2 P calcium imaging experiment with a drifting grating stimulation by the 270° monitor ring. **B:** Visual stimulation paradigm: in each stimulation cycle, after a gray screen, 8 directions appear in random order, comprising a static and drifting grating per direction. **C - E:** Calcium transients in response of the drifting grating stimulation and polar plots of the relative response levels of the identical longitudinally-tracked neuron in **C**: pre-induction (P-I), **D:** relapse (RL), and **E:** remission (RM). **F:** Direction selectivity indices (DSI) and **G:** Orientation selectivity indices (OSI) of all time points (median: **F**, DSI, P-I=0.33, RL=0.20, RM=0.25, \*\*\*\**p* _P-Iv.s.RL_<0.0001, \*\*\*\**p* _P-Iv.s.RM_<0.0001, *p* _RLv.s.RM_=0.50; **G,** OSI, pre-induction=0.33, relapse=0.23, remission=0.25, \*\*\**p* _P-Iv.s.RL_=0.0006, \*\**p* _P-Iv.s.RM_=0.0099, *p* _RLv.s.RM_=0.36, one-way ANOVA test). **H**: DSI and **I:** OSI of all activity levels of all time points (median: **H**, DSI, hypo-active=0, medium=0.33, hyper-active=0.16, \*\*\**p*_hypo-v.s.medium-_=0.0003, *p*_hypo-v.s.medium-_=0.26, \**p*_medium-v.s.hyper-_=0.045; **I**, OSI, hypo-active=0, normal=0.33, hyper-active=0.16, \**p*_hypo-v.s.medium-_=0.039, *p*_hypo-v.s.hyper-_=0.16, *p*_medium-v.s.hyper-_ =0.059, one-way ANOVA test). *n* = 5 animals and 267 neurons. **J-K**, activity sub-grouped DSI (J) and OSI (K) in all time points respectively. **L:** Experimental paradigm of visual discrimination task. Window shown white stimulus is correct choice, window shown one of gray screens (gray level from 10% (light gray) to 100% gray (black)) is false choice. **M:** Normalized correct-choice alteration upon presentation of different grayscale windows (100%, 50% and 10% are shown) and white windows at different time points. Control day 0 v.s. control day 21, EAE pre-induction v.s. EAE remission (control day 0 is comparable with EAE pre-induction; control day 21 is comparable with EAE remission, *p_cotrol- v.s. EAE @50%_ =0.0362). n = 16 animals (10x control, 6x EAE).

We next asked, whether - as in our assessment of the connectivity matrices upon spontaneous activity (Fig. 2) - the activity class of the respective neuron affects its visual tuning specificity. We found, that particularly the hyperactive neurons perform significantly worse than medium-active neurons (Fig. 7 H - I). This inability will most likely drastically impact the further upstream cortical regions, as both the ventral and dorsal stream, which eventually reach prefrontal circuits, originate in primary visual cortical areas (Gilbert and Wiesel, 1983, Ibbotson and Jung, 2020). Due to the imaging location in the visual cortex, we asked whether our observations might be triggered by an inflammation or demyelination process of the visual-afferent pathway. Therefore, we prepared the optic nerve of EAE mice in relapse and remission and determined CD4^+^ T cell, Mac-3 cell counts, and MBP staining (Suppl. Fig. 5). In EAE relapse we did observe a significant increase of CD4 cell infiltration in the optic nerve. This increase was reversed to almost healthy control levels in EAE remission. In addition, with regard to the number of activated macrophages we observed a significant increase of inflammation in relapse, ceding in remission. Myelination was assessed as fraction of myelinated area normalized to control (i.e. non-immunized animals). Here we observed a similar degree of myelination in all three groups. Together, while visual afferent pathways are affected by the disease process in relapse, these processes are limited to this disease stage, and vanish in remission, a stage in which we did observe a significant impairment of the encoding of visual afferents in the primary visual cortex.

Yet, to which extend are these impairments relevant for the behavioral discrimination ability of the animals? To evaluate the visual discrimination ability of SJL/J mice during remission, we conducted a visual discrimination task (Figure 7J, K). We trained the mice prior to EAE induction, to discriminate between a white screen and a darker screen (gray level from 10% to 100% gray (black), Fig. 7J). We then compared the discrimination ability three weeks after initial training. Both for very easy, and for very difficult discrimination tasks, no difference between EAE remission animals and controls could be observed. Notably, for a medium difficult discrimination task, EAE remission mice displayed a significant decrease in the discrimination ability between white and 50% gray. (Fig. 7K), indicating a distinct performance-impacting network maladaptation in visual cortical networks during remission.

### Decreased visual discrimination and object recognition in a cohort of stable and mildly affected RRMS patients

Finally, we asked, whether our finding of a performance-impacting network dysregulation particularly of visual cortical circuits in remission and in the absence of local inflammation is translationally relevant.

We thus investigated stable and only mildly (EDSS ≤ 3) affected human subjects with MS. We conducted a single-center cohort study with 20 stable RRMS patients without history of optic neuritis and unremarkable visual evoked potentials (VEPs), contrasted to 20 age/gender matched controls (Table 1). RRMS patients (mean age 35.2 ± 11.4) had a mean disease duration of 7.7 years and a mean time since last relapse of 5 years. The median EDSS was 1.25, indicating a rather low disease burden. Two RRMS patients were treatment naïve whereas 18 received a disease-modifying therapy.

**Table 1:** Demographic and clinical data of MS patients and healthy controls.

| Demographic and clinical data | Multiple sclerosis patients (n = 20) | Healthy controls (n=20) | P-value | Statistical test |
| --- | --- | --- | --- | --- |
| Sex (female/male) | 15/5 | 15/5 |  |  |
| Mean age at study (SD) [years] | 35.2± 11.4 | 34.35±9.9 | 0.86 | Mann-Whitney Test |
| Disease Course CIS/RRMS | 0/20 |  |  |  |
| Mean disease duration (SD) [years] | 7.7±5.7 |  |  |  |
| Mean time since last relapse (SD) [years] | 5.0±4.5 |  |  |  |
| Median EDSS score (Bierbrauer et al.) | 1.25 (0-2.5) |  |  |  |
| DMD (no / first-line / second-line) | 2/6/12 |  |  |  |
| Mean HADS-A score (SD) | 5.8±4.8 | 3.6±2.2 | 0.23 | Mann-Whitney Test |
| Mean HADS-D score (SD) | 3.6±4.9 | 1.4±1.8 | 0.26 | Mann-Whitney Test |
| Mean SDMT score (SD) | 58.1±8.5 | 60.4±9.9 | 0.43 | Unpaired t-test |
| Mean VLMT1-5 score (SD) | 58.2±9.1 | 61.6±5.9 | 0.18 | Unpaired t-test |
| Mean VLMT7 score (SD) | 12.6±2.2 | 13.8±1.5 | 0.059 | Mann-Whitney Test |
| Mean Munsell FM 100-Hue Test (SD) | 67.4±35.3 | 35.0±16.1 | ***<br>0.0009 | Unpaired t-test, Welch correction |
| Mean BORP (SD) | 28.9±2.1 | 30.3±1.2 | *<br>0.014 | Unpaired t-test, Welch correction |
| Mean sNfL (SD) [pg/ml] | 9.2±13.3 | 4.8±1.6 | 0.08 | Mann-Whitney Test |

For an optimized translational relevance, we subjected the patients to several visual discrimination tasks, which should reflect the ability of their visual circuits to adequately report subtleties of their visual environment (Fig. 8, Suppl. Fig. 8). To exclude any memory or cognitive impairments, we conducted a verbal learning and memory test (Suppl. Fig. 8A, VLMT 1-5, VLMT 7) and a concentration and decision-making test (Suppl. Fig. 8B, SDMT). The patients did perform these tests similar to the healthy controls, indicating that hippocampal and prefrontal circuits were not significantly affected at this stage. Neurofilament light chain (NfL, Suppl. Fig. 8C) was not significantly different between the two groups, indicating that MS patients showed no relevant subclinical neuroaxonal damage.

**Figure 8:**
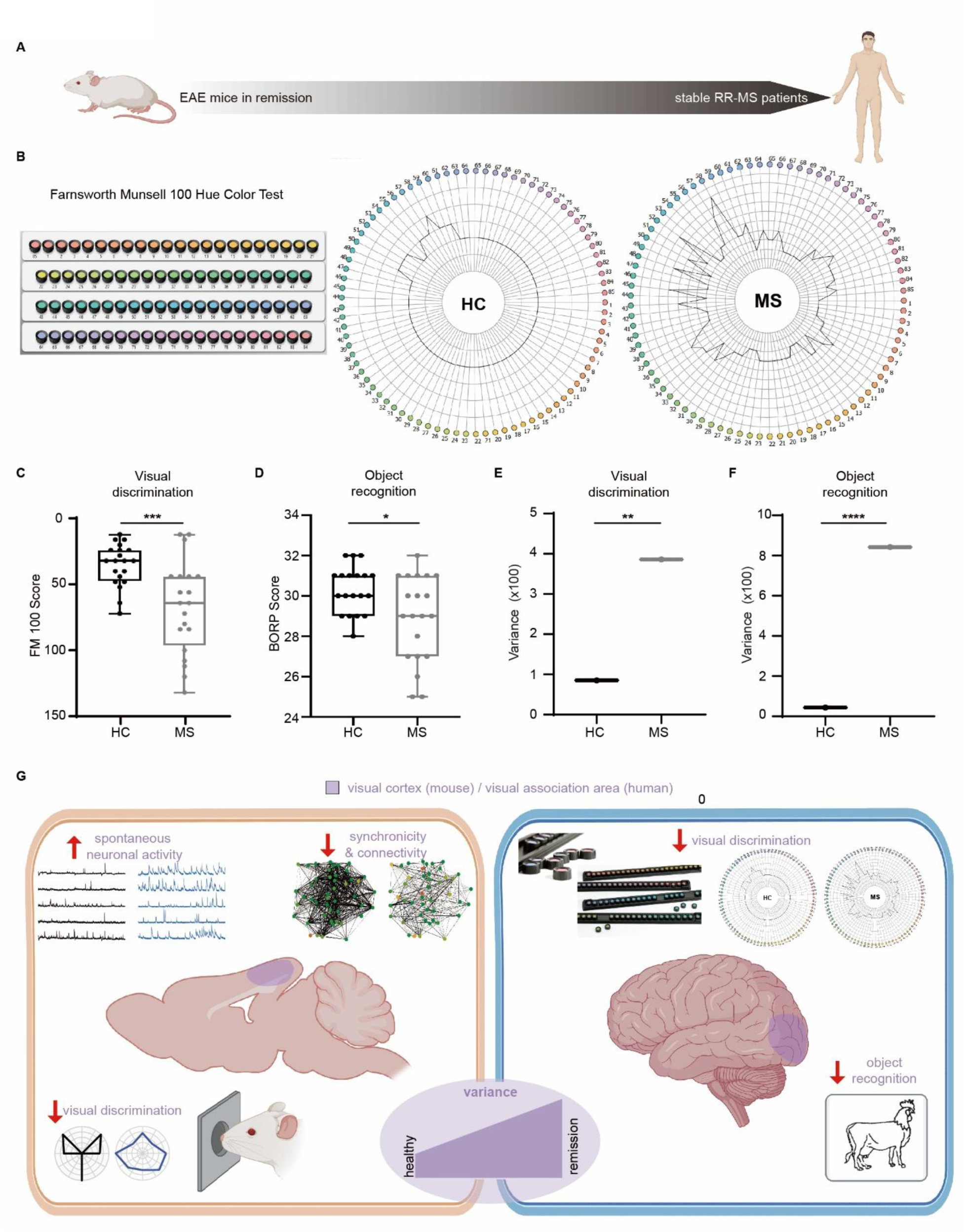
Impaired visual discrimination in RRMS patients. **A:** Translational study from mice to human. **B - F:** Stable and mildly affected RRMS patients and healthy controls underwent a test battery consisting of visual discrimination, object recognition measurement. **B, C:** Visual discrimination measured with Munsell FM 100-Hue Test revealed a significant worse performance in RRMS patients, mean ± sd, RRMS: 67.4±35.3; controls: 35.0±16.1, ***p<0.001, unpaired t-test with Welch correction. **D:** Object recognition (BORP) was decreased in RRMS, mean ± sd, RRMS: 28.9±2.1; controls: 30.3±1.2, *p=0.014, unpaired t-test with Welch correction. **E, F:** Variance of the rate of visual discrimination test (E) and object recognition test (F). Significant increase of variance in MS patients’ comparison to healthy control. Variance_FM100,HC_= 85.01, Variance_FM100,MS_ =386.01, ***P _FM100,HC v.s. MS_ =0.0025; Variance_BORP,HC_ =43.42, Variance_BORP,MS_ =841.94, **P_BORP, HC v.s. MS_ =3.86098E-07 (Levene’s test for the equality of variance). **G:** Overview of experimental data in visual cortex of animal and behavior experimental data in human. In animals, hyperactivity, decreased synchronicity and connectivity, and decreased visual discrimination ability were found in EAE remission phase. The variance of network change is significant higher in EAE remission mice when compare to healthy control. In human, decreased visual discrimination ability was found in stable and mildly affected RRMS patients when compare to healthy control.

Strikingly, RRMS patients performed significantly worse in visual discrimination as tested by the Munsell FM 100-Hue color test (Fig. 8B, C). In line, object recognition was also impaired in RRMS patients compared to healthy control subjects (Fig. 8D). Yet, while the mean performance was drastically and highly significantly reduced, the variance of the performers was significantly higher in RRMS patients (Fig. 8E, F). This indicates a high degree of variability in the extend of visual network dysfunction in our cohort, which is an observation often shared by clinicians particularly at early disease states (Engel and Zipp, 2022).

In sum, we report a novel and early brain state maladaptation (Fig. 8G), independent of the acute neuroinflammatory process, which might represent an early pathomechanism of progression independent from relapse activity in MS.

## DISCUSSION

Identifying early changes neuronal network function in neurological disorders is, potentially, essential for early diagnosis and treatment. Neuronal hyperactivity has been identified in many neurological disorders, including MS. The resulting brain state shifts may represent a unifying mechanism by which the bound network reacts upon internal challenges in the course of neurological disorders. Yet, particularly in MS, despite accumulating evidence on plasticity-mediated maladaptive network changes in animal models, both the directionality of network change – hypo- or hyperactivity – and the translational relevance of these initial findings remained unresolved. Here, by applying awake longitudinal 2-P calcium imaging in the visual cortex of a mouse model of RRMS (EAE mouse model), we find that initially, in the phase of active disease (1^st^ relapse) the overall rate of network activity remains unchanged. We unravel that both hypo- as well as hyperactivity represent a response by which the cortical network reacts to acute neuroinflammation, even with the first inflammatory attack on distant areas such as brainstem and spinal cord (Yang et al., 2022). In remission, in the absence of an acute inflammatory disease, only the hyperactive phenotype remains. The increased cortical network activity, accompanied by synaptic re-organization, is a new functional state leading to a significant deterioration of the local connectivity in the layer II / III of the visual cortex, based on spontaneous, ongoing recordings of neuronal activity. Cortical inflammation due to the virus injection or the EAE itself were not present. However, the brainstem, an area with prominent interaction between immune and neuronal cells in EAE (Siffrin et al., 2010), showed a significant reduction of optogenetically evoked calcium transients compared to healthy control animals. This indicate that the time-resolved shift towards neuronal hyperactivity and network dysfunction in the cortex came along with a reduced excitatory brainstem neurons response. This “silencing” even correlates with disease severity and progression. In line, transient neuronal silencing was already described in a focal EAE model with cytokine induced local cortical inflammation (Jafari et al., 2021b). There, immune cells are attracted to the cortex area where cytokines were injected. This is followed by a reduction in neuronal activity. The brainstem provides multiple sensory inputs to the cortex (Guillery and Sherman, 2002) and our observation in the brainstem might trigger functional changes in the far distant and not inflamed cortex. The upregulation of cortical inhibitory synapses on the other hand might represent a beginning compensatory mechanism. Electrophysiological recordings in the hippocampus also provided clear evidence of a modified network state during remission as compared to the pre-induction state, yet devoid of hyperactivity. Thus, hyperactivity might represent a putative link to subsequent activity-dependent neurodegeneration, as only in cortical hyperactive circuits, an increase in early cell death could be detected. Subjecting the animals to a classical visual stimulation paradigm – drifting gratings – we could conclude that the new brain state in remission is associated with a deterioration of the primary function of this local circuit: the specific representation of the outside world, here the accurate representation of the angle of the gratings. This finding indicates, that recordings of spontaneous ongoing activity are very well suited to predict and assess the performance of a local circuit for its dedicated task. It so seems, that in remission, impaired optogenetically evoked neuronal activity in the brainstem leads to even far distant network changes, e.g. also in the visual cortex. The cortical network reaches a new maladaptive state, characterized by hyperactive neurons and an impaired local circuit performance. And indeed, the deterioration in the encoding of visual afferents directly related in a distinct in ability of mice to perform in a medium-difficult discrimination task.

But, does this state persist, and does this finding bear any significance for human disease? In the SJL-EAE animal model, the remission phase is unfortunately temporally limited, as the remission is followed by additional relapses (Krishnamoorthy and Wekerle, 2009), and hence studies in animals reach their limit for assessing the stability of this new set point. Consequently, we now chose to investigate MS patients in a stable disease phase of remission, without any history of optical neuritis. Based on our findings in animal models, we would predict a rather subtle yet persistent network shift. Therefore, we chose a visual discrimination task which is rather difficult to perform. Indeed, it so seems that even in these early stages, the network needs to be challenged to reveal maladaptive network shifts (Bierbrauer et al., 2020). Stable and mildly affected RRMS patients performed worse in visual discrimination and object recognition compared to age- and sex-matched healthy controls, despite no history of optic neuritis and unremarkable visual evoked potentials. Notably, while the average performance was worse in patients, the variance in the performance was significantly increased in the patients, pointing towards a high degree of heterogeneity with regard to individual trajectories of network adaptation or maladaptation (Stroh et al., 2024). These distinct behavioral changes were not caused by impaired concentration, cognition (normal SDMT) (Benedict et al., 2017, Parmenter et al., 2007) or learning and memory (normal VLMT 1-5 and VLMT 7) (Helmstaedter and Durwen, 1990) in our cohort. Visuoperceptual impairment has been already described in MS patients (Vleugels et al., 2000). However, these MS patients had a longer disease duration, history of optic neuritis was not an exclusion criteria and most of the MS patients had a progressive disease course (Vleugels et al., 2000). We here investigated a stable mildly affected RRMS cohort (median EDSS is 1.25) with no functional affection of the visual tract. Thus, impairment in visual discrimination starts already early during disease and in the absence of any clinical affection of the visual tract. Another study, also including mainly older and progressed MS patients, revealed that visual object recognition is altered in cognitively impaired MS patients (Laatu et al., 2001). In our cohort, basic learning and concentration tests were unremarkable between MS and healthy control subjects. Therefore, cognitive deterioration did not explain the deteriorated performance in visual discrimination and object recognition in our cohort. A recent study detected impaired visual discrimination in older RRMS patients (mean age 43 years), but not in younger RRMS patients (mean age 37 years), although they used different tests (Caputi et al., 2017). Differences in inclusion and exclusion criteria (history of optic neuritis) might additionally explain the discrepancy to our study, where we find alteration of visual discrimination already early on. Network alterations due to altered neuronal activity might be the direct cause for disease symptoms in RRMS patients. Mapping disease symptoms to certain brain networks has recently become available and will help to understand and treat MS disease symptoms by a targeted, network-state-informed approach (Fox, 2018, Tetreault et al., 2020, Boes et al., 2015).

In the acute phase of RRMS, neuronal function even in distant nodes to the major lesion areas, e.g. brainstem or cerebrum, is severely affected. As brain networks need to retain a stable mode of functioning (Rosenthal et al., 2020, Stroh et al., 2024), they employ mechanisms of homeostatic plasticity to stabilize this new – maladaptive – set point of network function (Turrigiano, 2012).

This maladaptive brain state in remission has two negative consequences: first, the local network performance does not return to pre-relapse levels. A subtle yet significant decrease of visual performance levels is retained. Secondly, and maybe most importantly, a shift of brain state towards a state involving higher levels of hyperactive neurons may be associated with a decreased survival probability of these very neurons (Ellwardt et al., 2018). Our findings point to a highly dynamic, stage and disease-specific network re-organization in patients, as they exhibit a cortical network dysfunction reflected by decreased visual discrimination ability years after the last relapse.

The pathomechanisms behind progression occurring independent of relapse activity are not fully understood so far (Lublin et al., 2022, Kappos et al., 2020). Diffuse microglial activation, intrinsic progressing astrogliosis, failure of remyelination and neuronal loss include suspected mechanisms. We hypothesize that network alterations due to dysfunctional cortical activity may contribute to the occurrence of PIRA independent from direct inflammatory damage. In fact, there is already evidence that neuronal hyperactivity causes apoptosis in mice (Ellwardt et al., 2018) and that network alterations cause disability and cognitive impairment in MS (Jandric et al., 2021, Charalambous et al., 2020, Sjogard et al., 2021). We here report a direct link between cortical dysfunction in microcircuits in the absence of inflammation and impaired functional performance in EAE and early RRMS. Similar phenomena have been reported in other neurodegenerative diseases. In HD, the cortical network change is believed to be functionally correlated to its most severe neurodegeneration region, the striatum (Arnoux et al., 2018, Blumenstock and Dudanova, 2020). In AD, atrophy in cortical regions might associate with hippocampal hyperactivation, which might be an early indicator of AD-related neurodegeneration in a distributed network (Busche et al., 2012, Putcha et al., 2011). Counterbalancing this hyperactivity by treatment with the anticonvulsant drug levetiracetam might improve cognitive function in patients with mild cognitive impairment (Bakker et al., 2012). These data suggest that neural networks might undergo fundamental changes very early also in neuroimmunological disorders in a disease-transcending manner, in regions not yet affected by the underlying molecular and cellular pathophysiology. What do we learn from these concepts for network-based-therapies? Targeting maladaptive network dysregulations, particularly in phases of the disease in which no acute neuroinflammatory events occur has the potential to both prevent activity-dependent neurodegeneration and alleviate the subtle neuropsychiatric symptoms found in patients in remission. For making a network-centered therapy a reality, we need to further functional neuroimaging approaches in conjunction with a neuropsychiatric assessment, for an individualized and tailored intervention.

## Materials and Methods

### Experimental animals and EAE induction

Female 8–10-week-old SJL/J mice were acquired from Charles River Laboratory (Germany) or Janvier Labs (Bierbrauer et al.). All animals were group-housed at the Mouse Behavioral Unit, Johannes Gutenberg University Mainz and had ad-libitum access to food and water. All experimental procedures were performed in accordance with institutional animal welfare guidelines and were approved by the state of Rhineland-Palatinate, Germany. Randomization was carried out as follows: On arrival from company, each animal was assigned a random number and the experimental order was randomized. For SJL-EAE induction, mice were injected subcutaneously (Polman et al., 2011) with 200 µg PLP_139-151_ peptide (Hooke Laboratories) emulsified with complete Freund’s adjuvant (CFA, DIFCO) supplemented with 4 mg / mL Mycobacterium tuberculosis H37RA (DIFCO). Each mouse received one intraperitoneal (i. p.) injection (500 ng) of pertussis toxin at day 0 post immunization. EAE was scored clinically on a daily basis according to a 0 – 5 scale as follows: 0, no clinical symptoms; 1, tail paralysis; 2, impaired righting reflex and partial hind limb paralysis; 3, complete bilateral hind limb paralysis; 4 total paralysis of hind limbs and partial fore limb paralysis; and 5, dead. Due to obvious clinical symptoms the experimenter could not be blinded with regards to the stages of the animal during the experiment, but the data analysis was carried out with a standard analysis pipeline without bias.

### Virus injection, implantation of chronic window and holder

For longitudinal 2-P calcium imaging, implantation of the chronic window and holder was conducted while virus injection. Stock solutions of rAAV1/CamKIIa-GCaMP6f (3.67 x 10^11^ / ml, Penn Vector Core, University of Pennsylvania, PA, USA) was diluted with PBS to 1.84 x 10^10^ / ml and stored at 4 °C short before the injection. Mice anesthetized with isoflurane in O_2_ (Abbvie, Ludwigshafen, Germany) were placed in a stereotactic frame (Kopf Instruments, CA, USA) and warmed with a heating pad (World Precision Instruments, Sarasota, FL, USA). For chronic window preparations, the skull was exposed by cutting off the scalp and well cleaned for fixing the head-holder. A round craniotomy with maximum diameter of 2.5 mm was carefully conducted over primary visual cortex (V1, anteroposterior − 2.8 mm, mediolateral − 2.8 mm). Diluted viral solution was injected via a custom-made tip-loading system, including a pulled micro-pipette (Hirschmann Laborgeräte, Eberstadt, Germany), a syringe and plastic tubing, using manual pressure. The pulled micro-pipette was slowly inserted into the exposed brain and approximately 300 nl of the viral solution was injected with an injection speed around 0.2 μl / min stepwise from 600 μm depth, targeting layer V / VI, to 200 μm, targeting layer II / III. Before retraction, the micro-pipette remained in place for 5 - 10 minutes helping the dispersion of virus solution in brain tissue meanwhile preventing an outward flow. After the virus injection, the opening was closed with a circular cover slip (Electron Microscopy Sciences, Hatfield, PA, USA) with a diameter of 3 mm and sealed onto the skull with surgical glue. The head holder was fixed onto the exposed skull with UV-glue (Polytec, Waldbronn, Germany). For a more detailed protocol please see (Guimarães Backhaus et al., 2021).

For brainstem virus injections, skin and muscles were dissected and injection took place just below the cerebellum (Siffrin et al., 2010). Viral solutions of rAAV1/Syn-GCaMP6f (1.84 x 10^10^ / ml, Penn Vector Core, University of Pennsylvania, PA, USA) AAV-CamKII.C1V1.mcherry (2.00 x 10^11^ / ml, Penn Vector Core, University of Pennsylvania, PA, USA) were delivered (Fu et al., 2021) by a micro-pipette as described above.

### Two-photon (2-P) *in vivo* awake imaging

*In vivo* recordings were conducted using a custom-built 2-P microscope equipped with a resonance scanner (LaVision Biotec) and a Ti:Sapphire laser operating at 920 nm wavelength as light source (Coherent). A 40 × (NA = 0.8; Nikon) water immersion objective was used for imaging. Image acquisition was controlled by ImSpector Pro software (LaVision Biotec) at a frame rate of 30.8 Hz with a field of view of 277 x 277 μm^2^. Habituation of GCaMP6f injected mice with chronic window and holder were performed 5 days before the awake recording to adapt the awake head-fixing imaging system. Spontaneous 2-P calcium imaging was recorded for 15 min for each trial. Calcium transients were represented as relative changes in fluorescence (Δf / f).

### Visual stimulation by virtual reality

A drifting grating visual stimulation was presented in a virtual reality (Polman et al.) system consisting of a movement detectible air-float jet-ball system (PhenoSys, Berlin, Germany) and a 270° monitor system for delivering visual stimulus (Backhaus et al., 2022). The distance between the screens and eyes of the head-fixed mouse in the awake-behaving 2-P imaging system was approximately 50-60 cm. From this position, the screens covered the whole field of visual perception. The stimulation protocol was adapted from the Rochefort group (Rochefort et al., 2011). Stimuli were conducted per cycles. In each stimulation cycle, 5 seconds of gray screen will be displayed at the beginning (Fig. 3B), followed by 8 directions (0°, 45°, 90°, 135°, 180°, 225°, 270°, 315°), first static (5 s) then drifting (5 s) (Rochefort et al., 2011). In each visual stimulation experiment, 10 stimulation cycles were performed and the sequence of 8 directions in each cycle appeared randomly to prevent the possibility of adaptation of mice to the stimulus.

### Optogenetic stimulation of brainstem neurons

Recordings of the brainstem were performed in light anesthesia with isoflurane by inhalation after exposing the brainstem as described elsewhere (Siffrin et al., 2010). The breathing rate was maintained at > 120 breaths per minute during recording. Imaging in light-anesthetized animals was performed at disease onset and at relapse and compared to control non-EAE mice. For optogenetic stimulation of C1V1 neurons in the brainstem, a 552nm laser, coupled into our custom-built two-photon microscope was activated each 20s for a time period of 200ms. This was repeated for 20 cycles. The PMT shutters were closed while stimulation in order to avoid imaging oversaturation.

### Analysis of spontaneous activity

Initial image processing was carried out using custom-written scripts in MATLAB (MathWorks Corporation, Natick, MA). A semiautomatic algorithm was used to detect cell outlines, which were subsequently confirmed by visual inspection. All pixels within the cell-based regions of interest (ROIs) were averaged to yield a time course (Δf / f) for each neuron, which was further analyzed by Igor software (Wave-Metrics). For more details please check our published protocol (Guimarães Backhaus et al., 2021).

#### Categorization of hypoactive, medium-active and hyperactive neurons

Cell activities described by the number of calcium transients per minute were divided into three categories: hypoactive (green), medium-active (yellow) and hyperactive (red). The categorization was based on cell transients in the pre-induction phase, where the 25% and the 98% of this phase were used as thresholds between the hypoactive, medium-active and hyperactive category for all three disease states: pre-induction (P-I), relapse (RL) and remission (RM). For the global transition of calcium transients, the individual transition distributions were concatenated. The Sankey plot for transition diagrams was created in R using the R-package “easyalluvial”.

### Computation of network correlation

#### Synchronicity

The peak positions of ROIs of each cell were used to determine the activity intervals for each ROI. This was done by defining the onset and offset of each peak as the first frame to the left and right of the peak frame, respectively, that had a df / f intensity < 0.5 * peak intensity. These intervals between onset and offset were defined as activity intervals. Peaks were considered from the earliest to the latest peak, and peaks that were already part of an activity interval were not considered in the subsequent estimation of onset/offset but were instead considered part of the existing activity interval. The computation of Δf / f traces for this analysis was adapted from the Allen Brain SDK 2.14 as a windowed median filter detrending with using a long filter of 5401 frames and a short filter of 101 frames (Allen Institute for Brain Science, 2022, Software Development Kit [2.14]. Available from https://allensdk.readthedocs.io/en/latest/).

This binary activity matrix of all ROIs was then used to create pairwise correlations between each pair of ROI in each state and is depicted in a heatmap as synchronicity for every disease state.

#### Connectivity

In order to visualize the connectivity between every single neuron, the pairwise ROI correlation within timeframes in which at least one ROI showed activity was used to construct a network graph. This network consists of nodes for every neuron. The nodes position is based on its x and y position while the edges between these nodes that represent the pairwise correlation by the edge’s thickness. Consequently, edges between ROIs with a pairwise correlation ≤ 0 (Tchumatchenko et al., 2010) are not visible. The nodes’ color further represents the related ROI’s activity, making it possible to detect connectivity and activity patterns at the same time.

### Electrophysiological recordings and analysis

Prior to electrophysiological recordings, mice received a subcutaneous injection of Rimadyl® at 4mg/ kg bodyweight. After 30 minutes, mice were deeply anesthetized with isoflurane and placed in a stereotactic frame. A linear incision was done in order to expose the skull and H_2_O_2_ was applied to remove connective tissues covering the skull. To generate a location for the ground wire, a first craniotomy was performed at the contralateral hemisphere. For the electrode, a second craniotomy was drilled at the following coordinates: anteroposterior, −2.7 mm; lateral, −2.2 mm; dorsoventral. The exposed brain and skull were covered with saline. The probe (NeuroNexus A1x16-3mm-50-177 and A1x16–3mm-100-177) was moved to the coordinates mentioned above and lowered 2 mm below the brain surface to target the hippocampus. The ground wire was placed between skull and brain in the craniotomy on the contralateral hemisphere. After adjusting the anesthesia to achieve a persistent brain state, the recording was started with a sampling rate of 20 kHz.

Recordings from 16 control and 17 EAE mice were analyzed. Raw data was extracted with Matlab using the Spike2 Matlab SON Interface (Cambridge Electronic Design) and saved as a binary file to conduct spike detection and sorting with Mountainsort 5 in SpikeInterface (https://spikeinterface.readthedocs.io/en/stable/). After spike sorting, units were curated by excluding all units with an ISI violation ratio parameter bigger than 1 and a total spike rate over the whole recording smaller than 0.2 Hz (282 remaining units out of 394 detected units, 160 in control and 122 in the EAE group). The first and last ten seconds of each recording were excluded from the analysis to avoid potential settling artifacts. Average unit spike rates were calculated as the number of spikes each unit fired across the total recording time. For the analysis of activity phases, spike occurrences across all units were combined for each animal and the recording time divided into one second phases, for which the spike rate was calculated and normalized by the number of included units for each animal.

### Analysis of visual stimulation data

Functional calcium traces of each defined cell (Tetreault et al.) during the drifting grating stimulation section were subjected to a peak detection algorithm as described above, resulting in a binarized trace, including the timepoints of putative action-potential-related calcium transient peaks (Guimarães Backhaus et al., 2021). The binarized traces of all identified neurons were overlayed with the time course of the stimulus representations. Both traces were recorded by dedicated AD-converter (Bierbrauer et al.) and software (Spike 2) to prevent temporal jitter. Based on calcium responses correlated to the individual visual stimulation sequences, polar plot was generated. According to the polar plot, first, the preferred direction θ_pref_ was defined as the orientation of the presented grating stimulus direction that elicited the most calcium transients during the measurement. If more than one grating stimulus direction elicited the same amount of calcium transients, θ_pref_ was defined as the direction where the highest amplitude of a calcium transient was measured. Then orthogonal responses θ_orth_ were calculated. In the end the orientation selectivity index (OSI) and direction selectivity index (DSI) could be calculated as:

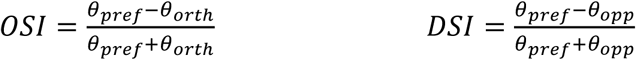

### Immunofluorescence staining, imaging, and quantification

For immunofluorescence staining, animals were perfused with PBS and 4% PFA and isolated brains and/or optic nerves were fixated in 4% PFA overnight. Brains and optic nerves were cut into 40 µm or 10 µm sections respectively. Staining was performed for CD4^+^ T cells (mouse anti-CD4-AF647, 1:150, 4010845, BD Biosciences), microglia (rabbit anti-Iba-1, 1:500, LER0547, Wako), activated macrophages (rat anti-CD107b [mac-3], 1:500, 2244706, BD Biosciences), caspase-3 (rabbit anti-caspase-3, 1:750, 47, Cell Signaling) and MBP (rat anti-MBP, 1:100, GR3360328-2, Abcam).

Overnight staining with the mentioned primary antibodies were followed by incubation with following secondary antibodies for 2 h: goat-anti rabbit Alexa Fluor 568 (1:1000; 2273773; Invitrogen) goat anti-rat Alexa Fluor 568 (1:1000; 2217022; Invitrogen); goat anti-rat Alexa Fluor 488 (1:1000; 2048174; Invitrogen); goat-anti rabbit Alexa Fluor 647 (1:1000; 2299231; Invitrogen). All sections were counterstained with DAPI (D1306, Thermo Fisher Scientific, 1:1000) before being transferred and embedded onto object plates.

Sections for immunohistology were analyzed using a Keyence epifluorescence microscope (Keyence, BZ-X700 Series), and mosaic scans with a 20× objective (Leica, NA 0.5) were performed. For assessing cell numbers per area and co-localization, cells were identified by semi-automated quantification software (ImageJ). The quantification of the immunofluorescence signal of MBP was performed with identical acquisition settings for each slice. No post-processing was performed. The percentage of myelinated area was calculated using the threshold tool with ImageJ as described elsewhere (Govier-Cole et al., 2019).

To determine the number of pre- and postsynaptic clusters and the number of synapses in the V1 cortical layers, 100 µm sections were cut by a vibratome. For each condition, 3 slices of 3 animals were analyzed. Layers II/III, layer IV and layers V/VI were detected in each slice using DAPI staining. The following antibodies and concentrations were used to analyze inhibitory and excitatory synapses: DAPI (D1306 Thermo-Fisher Scientific, 1:1000), anti-VGlut1 (guinea pig, 135304, Synaptic Systems 1:250), anti-Homer1 (chicken, 160006, Synaptic Systems, 1:250), anti-VGat (guinea pig, 131004, Synaptic Systems, 1:250), anti-Gephyrin (rabbit, 147018, Synaptic Systems, 1:250) with the respective secondary antibodies Alexa647-conjugated anti-guinea pig (goat, A-21450, Thermo-Fisher Scientific; 1:1000), Alexa488-conjugated anti-chicken (goat, A-32931, Thermo-Fisher Scientific; 1:1000), Alexa488-conjugated anti-rabbit (goat, A-32731, Thermo-Fisher Scientific; 1:1000). The slices were then mounted with ProLong Gold Antifade (P36930, Thermo-Fisher Scientific) and imaged with a confocal microscope (Leica TCS SP8, 63x objective, NA 1.3). To determine the number of pre- and postsynaptic clusters and the number of synapses, the synapse counter plugin of Fiji (Schindelin et al. 2012) was used as described previously (Dzyubenko et al. 2016) with the following parameters: rolling ball radius 1, maximum filter radius 2, threshold adjustment Otsu, pre- and postsynaptic particle size 12 voxels, maximum pre- and postsynaptic particle size 750 voxels.

### RNAscope

Adult SJL/J mice were perfused with PBS and 4% PFA. Brains were cut into 20-µm thick sagittal sections and mounted on Superfrost Plus Gold slides (Menzel). RNAscope in situ hybridization was performed using the Multiplex Fluorescent Detection Kit v2 (323110; Advanced Cell Diagnostics). Briefly, sections were dehydrated through a series of increasing ethanol concentrations (50%, 70%, and 2× 100%). After a 10-minute incubation with hydrogen peroxide at room temperature and antigen retrieval in 1× RNAscope target retrieval solution (ACD) for 5 minutes at 99°C, sections were treated with Protease III (ACD) for 30 minutes at 40°C. The following target probes were used in this study (all from ACD): Mm-Ripk1 (464511), Mm-Nefh-C4 (443671), and Mm-caspase 3 (441781). Hybridization was carried out for 2 hours at 40°C. Signal amplification and detection were performed according to the manufacturer’s instructions using TSA Vivid fluorophore 570 (1:500) and TSA Vivid fluorophore 520 (1:500). After counterstaining with DAPI for 30 seconds, sections were mounted with ProLong Gold Antifade reagent.

### RNAscope data analysis and quantification

Sections were analyzed using a Leica confocal microscope (Leica TCS SP8, DM-6000CS) with 40x objective and identical settings for each slide.

Quantification of target signal (RIPK1 and Caspase 3) was performed in two different manners. In brief, 10 randomly picked NEFH positive cells were selected in for each condition. For puncta per cell analysis, each signal was counted individually. Signal intensity scores were defined according to ACB criteria (see table 2).

**Table 2:** Signal intensity scores.

| Score | Criteria |
| --- | --- |
| 0 | No staining or <1 dot/10 cells |
| 1 | 1-3 dots/cell |
| 2 | 4-9 dots/cell, no or very few dot clusters |
| 3 | 10-15 dots/cell and/ or <10% dots in clusters |
| 4 | >15 dots/cell and/ or >10% dots are in clusters |

### Mouse visual discrimination task

In order to keep animals motivated to perform the task, daily water of animals in their home cage was added with 2% citric acid (CA) 5 days prior to experiment start, allowing animals more control over their water intake without interfering with behavioral performance (Urai et al., 2021). The experiment was performed in an isolated operant chamber with a touchscreen including two side by side response windows and a magazine with water dispenser on the opposite side (K-Limbic, Med Associates Inc., St. Albans, Vermont, Fig. 3I). The experiment consisted of following steps: 1) “reward training”: all animals were habituated individually for associating illumination combined sugar water reward in magazine (whenever sugar water was given, there will be illumination in the magazine). Completion conditions: drink 60 times of 10% sugar water giving in magazine within 30 min. for three consecutive days; 2) “touch screen training”: in this step, one response windows will be shown on the touch screen with white signal (correct stimulus). Animals needed to first trigger the sugar water reward in the magazine by nose poke the response window, then drink the reward from the magazine to finish one reward loop. Completion conditions: trigger 60 rewards loops within 30 minutes for 3 consecutive days; 3) “discrimination training”: in this step, two response windows will be shown on the touch screen, pairwise stimulus (black and white) were shown in these two response windows randomly. Animals needed to select and touch the correct stimulus (white) window to trigger a sugar water reward and drink it. Completion conditions: trigger more than 60 rewards within 30 minutes for 3 consecutive days; 4) “visual discrimination experiment (baseline)”: finally, well-trained healthy animals were ready for visual discrimination task. In this step, stimulus were conducted per cycles. In each stimulation cycle, correct stimulus (white) and one of 6 randomized appeared false stimuli (gray level of 10%, 30%, 50%, 70%, 90%, 100%) were pairwise shown randomly in two response windows. Animals needed to distinguish and nose poke the correct stimulus (white) to get the reward; 10 cycles was performed resulting 60x pair stimulus in total. Experiment ended after 10 cycles of stimulus. Afterwards, animals were randomly divided into two groups: one group was induced with EAE the following day (EAE group), and the other group was kept as healthy control group. All animals were maintained back to normal condition (normal drinking water) after the 4^th^ step until 5 days prior to remission phase, all drinking water was again replaced with 2% CA water; 5) “visual discrimination experiment (remission)”: when EAE group reached remission phase, the visual discrimination experiment described in 4) was repeated on both EAE and healthy control animals.

### Human cohort to test visual discrimination

Eligible RRMS patients were identified in our neurological outpatient clinic (Mainz, Germany). The local ethics committee approved the protocol. Twenty RRMS patients and 20 healthy controls have been included. Baseline characteristics can be appreciated in Table 1.

<u>Inclusion criteria</u> were as follows: age ≥ 18 and ≤ 65 years; EDSS ≤ 3; no relapse or cortisone treatment in the last 4 weeks.

<u>Exclusion criteria</u> were the following: symptoms of optic neuritis in the history; pathological visual evoked potentials (VEP); primary or secondary progressive disease course.

The <u>test battery</u> for all patients and controls consisted of the Farnsworth-Munsell (Wiendl et al.) 100-Hue test for visual color discrimination (Vleugels et al., 2000, Farnsworth, 1957), object recognition by the Birmingham object recognition battery (BORP, Test 10A) (Riddoch and Humphreys, 1993), anxiety and depression measured by the hospital anxiety depression scale questionnaire (HADS-A and D) (Zigmond and Snaith, 1983), concentration, cognition and decision-making by the symbol digit modalities test (SDMT) (Benedict et al., 2017, Parmenter et al., 2007), learning and memory by the auditory verbal learning and memory test (German version VLMT 1-5 and VLMT 7) (Helmstaedter and Durwen, 1990). Additionally, one blood sample was collected in order to determine serum neurofilament light chain (sNfL) concentration.

### Statistical Analysis

Statistical analyses were performed using GraphPad Prism 9 (GraphPad Software, Inc.), SPSS (Version 28, IBM SPSS Statistics) and MATLAB. All data sets were subjected to the Shapiro-Wilk normality test. For comparison of more than two normally distributed groups, the presence of significant differences was first evaluated using ordinary one-way ANOVA, followed by Dunnett’s multiple comparisons test against the control values. For comparison of more than two not normally distributed groups, the presence of significant differences was first evaluated using ordinary Kruskal-Wallis test, followed by Dunn’s multiple comparisons test against the control values. Variances statistical analysis was evaluated by Levene’s test. Quantification results were shown as a whisker plot (10-90 percentile, median). Chi-Square test was applied to investigate differences in frequency of spontaneous or optogenetically evoked neuronal activity.

To compare the two groups in the human cohort, normality was tested as stated above and either two-sided unpaired students t-test or Mann-Whitney test were applied. When variances were significant different, a Welch-correction was applied.

## List of Supplementary Materials

Supplementary Figure 1: Locomotion of each individual experimental mouse during awake 2-P calcium recording does not impact neuronal activity in V1.

Supplementary Figure 2: Spontaneous activity pattern remains stable in control mice over time.

Supplementary Figure 3: Individual animal data on neuronal activity

Supplementary Figure 4. No signs of inflammation in the cortex following virus injection and during EAE.

Supplementary Figure 5. No significant inflammation or demyelination of optic nerve in EAE remission.

Supplementary Figure 6. Number of excitatory and inhibitory synapses in Layer II + III, IV and V + VI.

Supplementary Figure 7. Neuronal cell death staining reveals no widespread neurodegeneration in EAE.

Supplementary Figure 8: Test battery for patients and controls beside visual discrimination and BORP.

## Data and materials availability

All data associated with study will be made available upon request.

The code for analysis can be found at: https://github.com/Strohlab.

## Author contributions

Study design: AS., FZ., SB., and EE.

Calcium imaging performing: TF.

Calcium imaging analysis: TF, NR. and AW

Animal behaviour experiment performing: TF, KE, and KR.

Animal behaviour experiment data collection and analysis: TF.

Electrophysiology experiment performing: KE, TF and WF.

Electrophysiology experiment analysis: TdS, ML, WO, TF, KE. and WF.

Immunostaining: MS, ME, MR, MS and KE.

Immunostaining analysis: EE, MS, ME, MJS.

Brain stem data collection and analysis: EE.

RNA scope experiment and analysis: ANS.

Human patient data collection: EE.

Statistical analysis: TH.

Writing: AS, TF, FZ, EE, LK and SB.

## Abbreviations

EAE: Experimental Autoimmune Encephalitis
FJC: Fluoro Jade C
MS: Multiple Sclerosis
NfL: Neurofilament Light chain
PIRA: Progression Independent from Relapse Activity
RIPK1: Receptor-interacting serine/threonine protein kinase 1
RRMS: Relapsing Remitting Multiple Sclerosis
VLMT: Verbal Learning and Memory Test

## Supplementary Materials

**Supplementary Figure 1:**
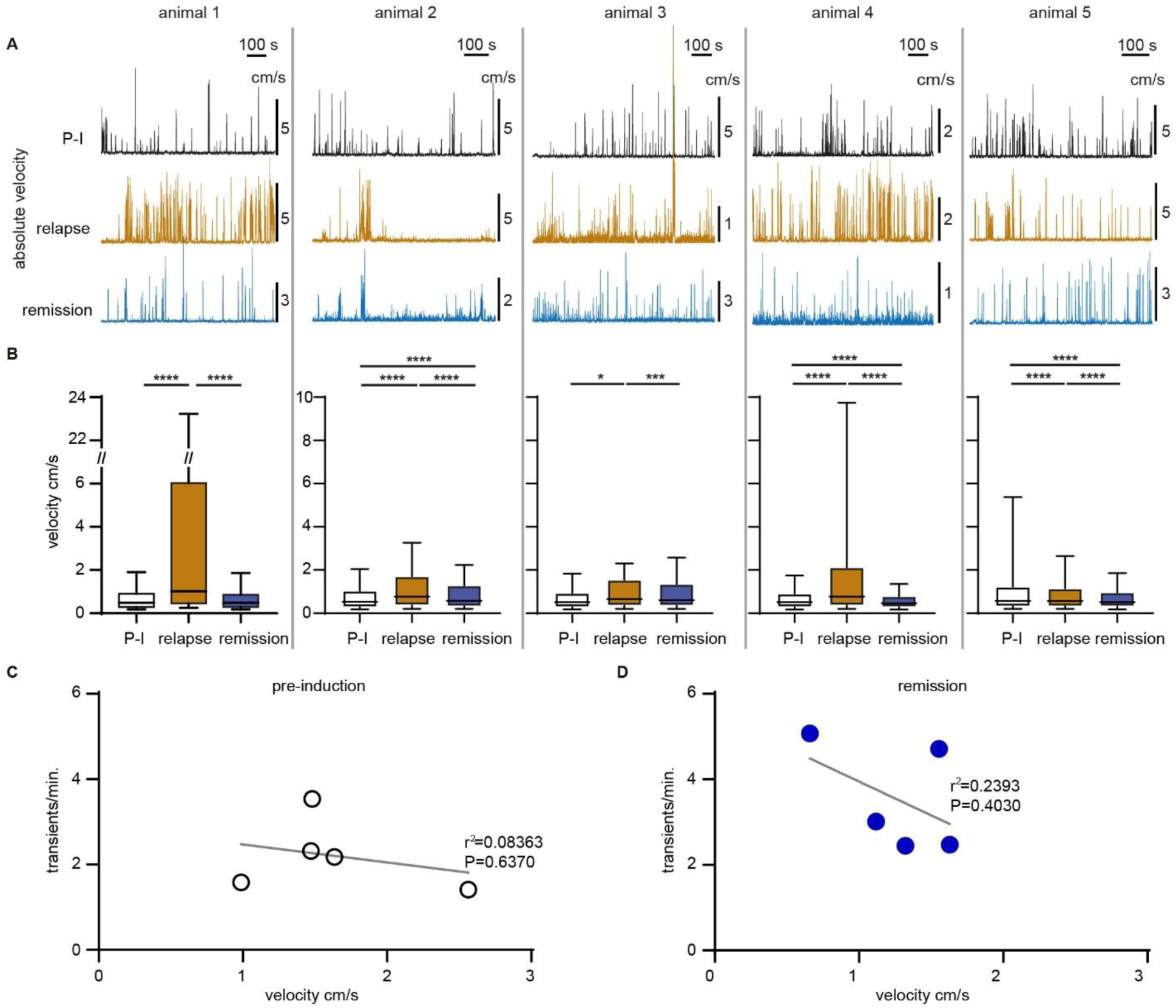
Locomotion of each individual experimental mouse during awake 2-P calcium recording does not impact neuronal activity in V1. **A:** Velocity traces of individual experimental mouse (animal 1 - 5) of awake 2-P calcium imaging in three time points: pre-induction (P-I), relapse and remission. **B**: Quantification of velocities in the same mice in all three time points (median: *animal 1*: pre-induction (P-I) 0.0016, relapse (RL) 0.06, remission (RM) 0.016, \*\*\*\**p* _P-Iv.s.RL_<0.0001, \*\*\*\**p* _RLv.s.RM_<0.0001, *p* _P-Iv.s.RM_=0.9984; *animal 2*: pre-induction (P-I) 0.0053, relapse (RL) 0.0077, remission (RM) 0.0058, \*\*\*\**p* _P-I v.s. RL_< 0.0001, \*\*\*\**p* _P-I v.s. RM_< 0.0001, \*\*\*\**p* _RL v.s. RM_< 0.0001; *animal 3*: pre-induction 0.0051, relapse 0.0065, remission 0.0061, \**p* _P-I v.s. RL_= 0.016, *p* _P-I v.s. RM_= 0.35, \*\*\**p* _RL v.s. RM_= 0.0001; *animal 4*: pre-induction 0.0051, relapse 0.0077, remission 0.0046, \*\*\*\**p* _P-I v.s. RL_< 0.0001, *p* _P-I v.s. RM_< 0.0001, *p* _RL v.s. RM_< 0.0001; *animal 5*: pre-induction 0.0057, relapse 0.0057, remission 0.0051, \*\*\*\**p* _P-I v.s. RL_< 0.0001, \*\*\*\**p* _P-I v.s. RM_< 0.0001, *p* _RL v.s. RM_< 0.0001; one-way ANOVA test). **C:** Similar pattern of relationship between the calcium transient rate to locomotion speed in both pre-induction and remission phase. Each point represents one animal.

**Supplementary Figure 2:**
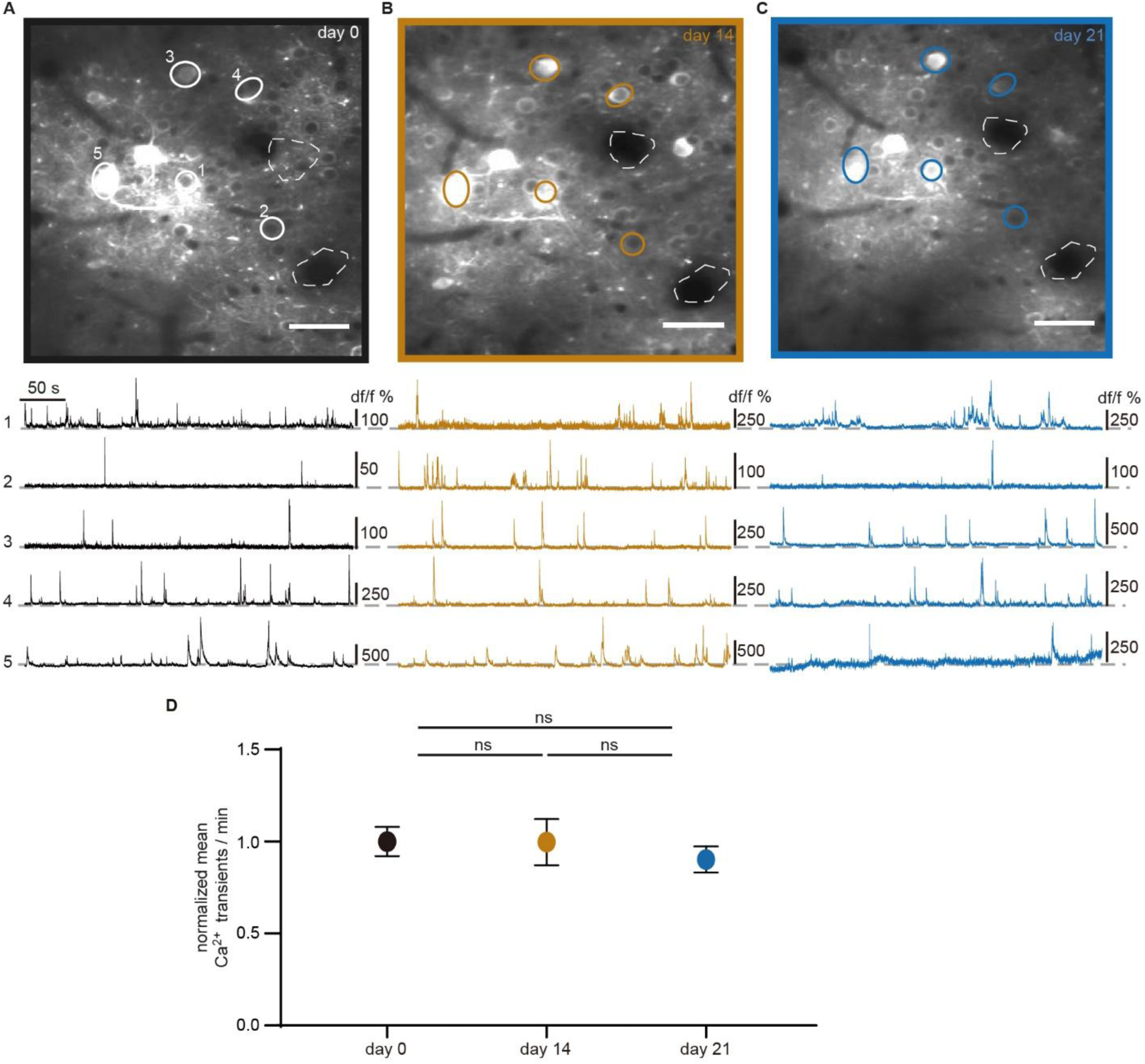
Spontaneous activity pattern remains stable in control mice over time. **A - C**: 2-P imaging (upper) and corresponding calcium transients (below) of identical neurons in layer II + III of primary visual cortex of an awake behaving mouse at three different time points of healthy mice, corresponding to three different time points of an EAE mouse: **A** Day 0 = EAE pre-induction (black), **B** Day 14 = EAE relapse (brown), **C** Day 21 = EAE remission (blue). Scale bar =100µm. **D**: normalized calcium transient rate (firing rate) per minute in day 0, day 14 and day 21. No significant change was found. **E:** Average calcium transient rates of each EAE animal (circles) at different disease stage (P-I: pre-induction, relapse and remission). Dashed line represents the mean value of all 5 animals.

**Supplementary Figure 3:**
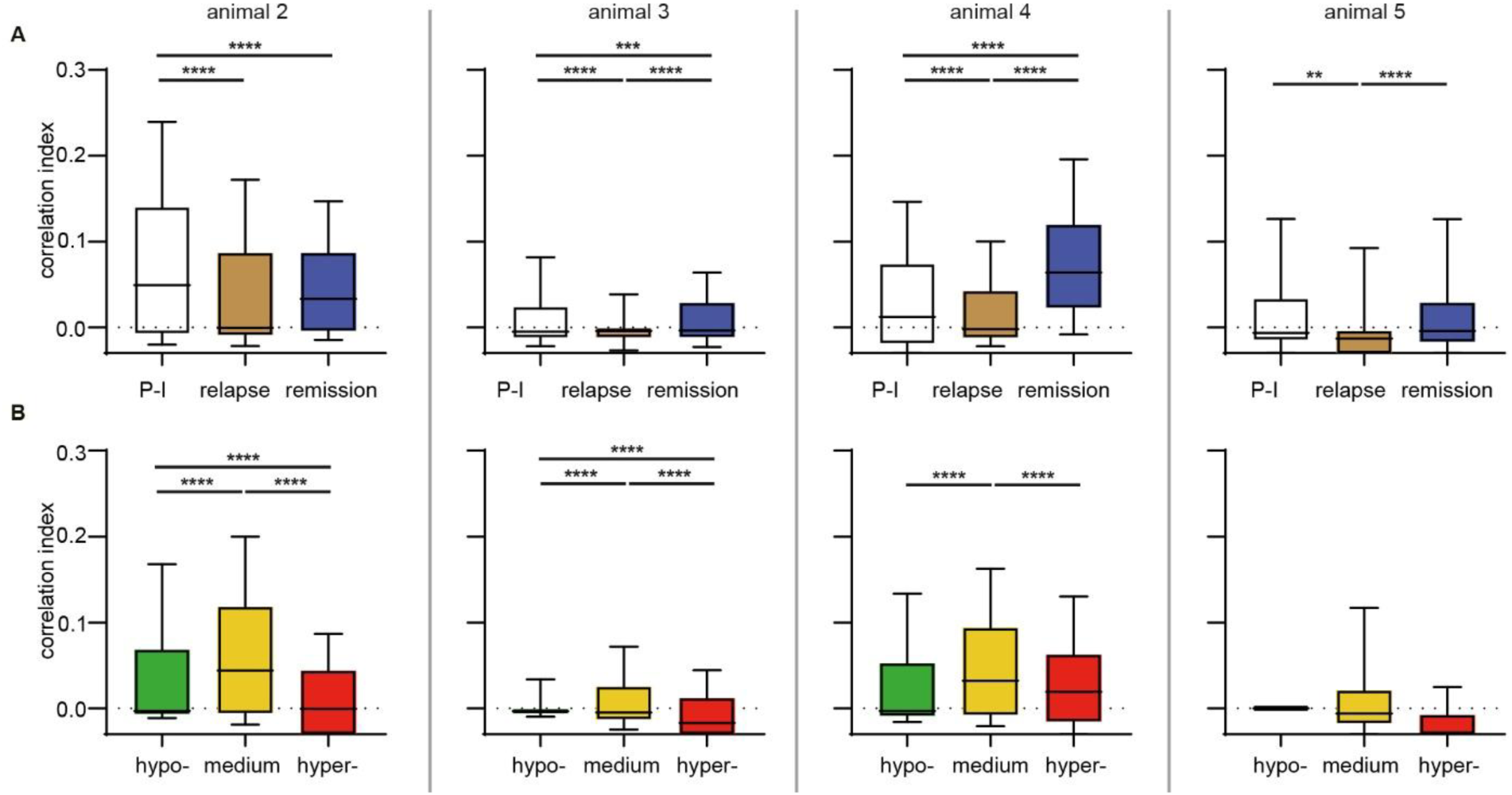
Individual animal data on neuronal activity. **A**: Correlation indices of individual mouse between three time points: pre-induction (P-I), relapse and remission. Median: *animal 2*: pre-induction 0.049, relapse −0.00053, remission 0.033, \*\*\*\*\**p* _P-I v.s. RL_< 0.0001, \*\*\*\**p* _P-I v.s. RM_< 0.0001, *p* _RL v.s. RM_= 0.31; *animal 3:* pre-induction −0.0047, relapse −0.0040, remission −0.0034, \*\*\*\*\**p* _P-I v.s. RL_< 0.0001, \*\*\**p* _P-I v.s. RM_= 0.0009, \*\*\*\**p* _RL v.s. RM_< 0.0001; *animal* 4: pre-induction 0.013, relapse −0.0019, remission 0.064, \*\*\*\*\**p* _P-I v.s. RL_< 0.0001, \*\*\*\**p* _P-I v.s. RM_< 0.0001, \*\*\*\**p* _RL v.s. RM_< 0.0001; *animal 5:* pre-induction −0.0063, relapse −0.013, remission −0.0040, \*\**p* _P-I v.s. RL_= 0.006, *p* _P-I v.s. RM_= 0.70, \*\*\*\**p* _RL v.s. RM_< 0.0001). **B:** Correlation indices of individual mouse between the activities levels of all time points combined. Median: *animal 2*: hypo-active −0.0028, medium-active 0.044, hyper-active −0.00039, \*\*\*\*\**p* _hypo- v.s. medium-_< 0.0001, \*\*\*\**p* _hypo- v.s. hyper-_< 0.0001, \*\*\*\**p* _medium- v.s. hyper-_< 0.0001; *animal 3*: hypo-active −0.0031, medium-active −0.0049, hyper-active −0.017, \*\*\*\**p* _hypo- v.s. medium-_< 0.0001, \*\*\*\**p* _hypo- v.s. hyper-_< 0.0001, \*\*\*\**p* _medium- v.s. hyper-_< 0.0001; *animal 4*: hypo-active −0.0030, medium-active 0.032, hyper-active 0.019, \*\*\*\**p* _hypo- v.s. medium-_< 0.0001, *p* _hypo- v.s. hyper-_= 0.12, \*\*\*\**p* _medium- v.s. hyper-_< 0.0001; *animal 5*: hypo-active 0, medium-active −0.0058, hyper-active −0.053, *p* _hypo- v.s. medium-_= 0.92, *p* _hypo- v.s. hyper-_= 0.33, \*\*\*\**p* _medium- v.s. hyper-_< 0.0001). *n* = 4 animals and 221 neurons.

**Supplementary Figure 4.**
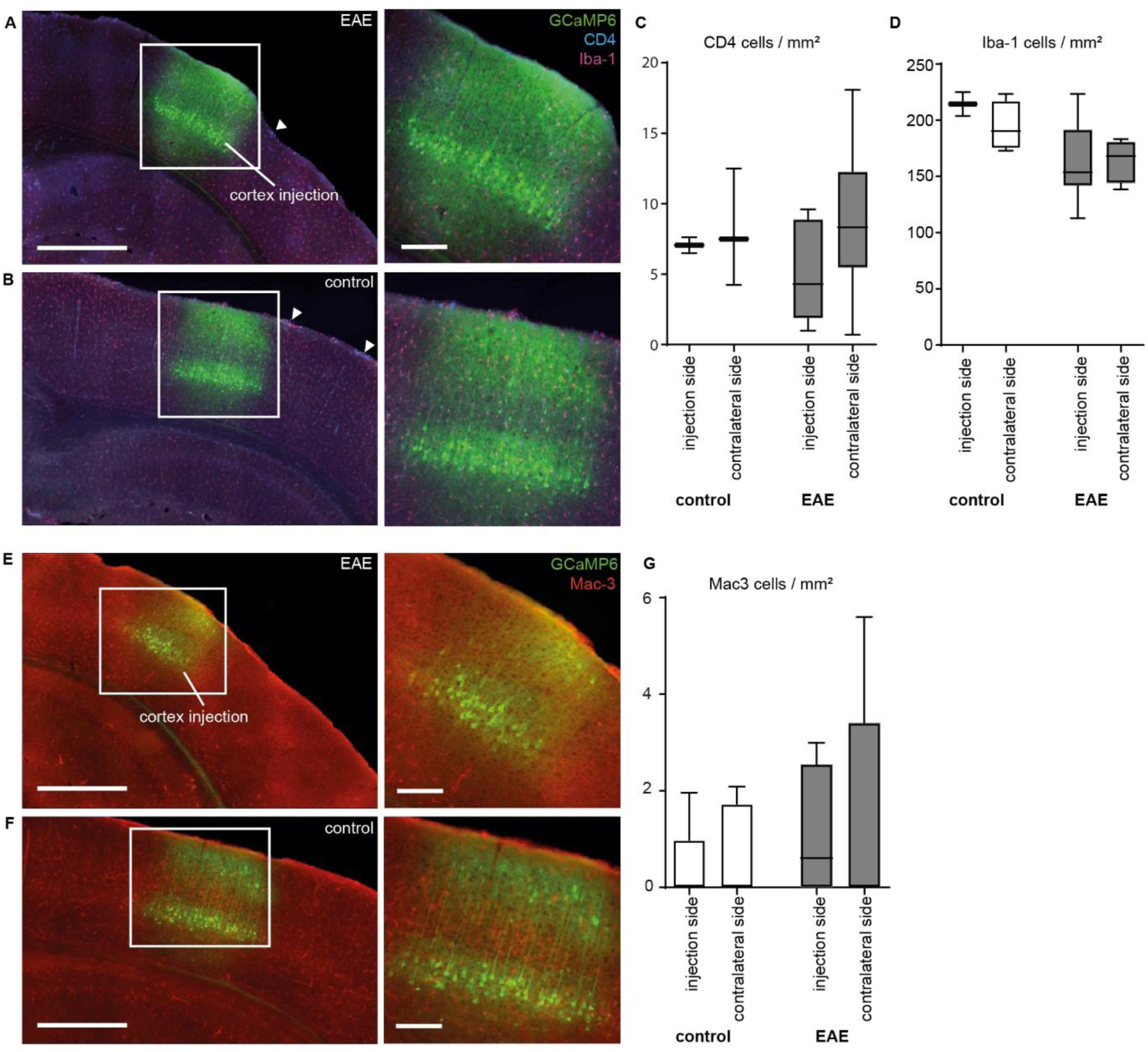
No signs of inflammation in the cortex following virus injection and during EAE. **A:** Expression of GCaMP6 (green) in V1 cortex and staining for CD4^+^ T cells (blue) and Iba-1 positive microglia cells (magenta) in EAE remission. **B:** Expression of GCaMP6 in V1 cortex and staining for CD4^+^ T cells and Iba-1 positive microglia cells in control mice. **C:** Only a few CD4 cells could be visualized at the site of GCaMP6 injection and non-injected contralateral cortex half with no significant difference between EAE and control mice: mean ± SD, EAE: 5.01 ± 3.47 CD4 cells / mm² ipsilateral and 8.51 ± 5.52 CD4 cells / mm² contralateral, *n*= 3 animals and 2-7 slices, *p* > 0.05. **D:** Quantification of Iba-1 cells revealed no differences under EAE condition at the site of GCaMP6 injection and non-injected contralateral cortex half compared to non-immunized mice. Mean ± SD, EAE: 162.67 ± 36.89 Iba-1 cells / mm² ipsilateral and 165.47 ± 18.16 Iba-1 cells / mm² contralateral, *p* > 0.05. **E:** Expression of GCaMP6 (green) in V1 cortex and Mac-3 positive activated macrophages (red) during EAE remission. **F:** Expression of GCaMP6 (green) in V1 cortex and Mac-3 positive activated macrophages (red) in control mice. **G:** Almost no Mac-3 positive cells were detected in EAE and control conditions as well as at the site of GCaMP6 injection and the contralateral site: mean ± SD, EAE: 1.05 ± 1.41 Mac-3 cells / mm² ipsilateral and 1.36 ± 2.42 Mac-3 cells / mm² contralateral, *p* > 0.05. P values were determined using one-way ANOVA test. Scale bars= overview images 300 µm and inserts 150 µm. *n* = 2-4 animals and 2-4 slices per group.

**Supplementary Figure 5.**
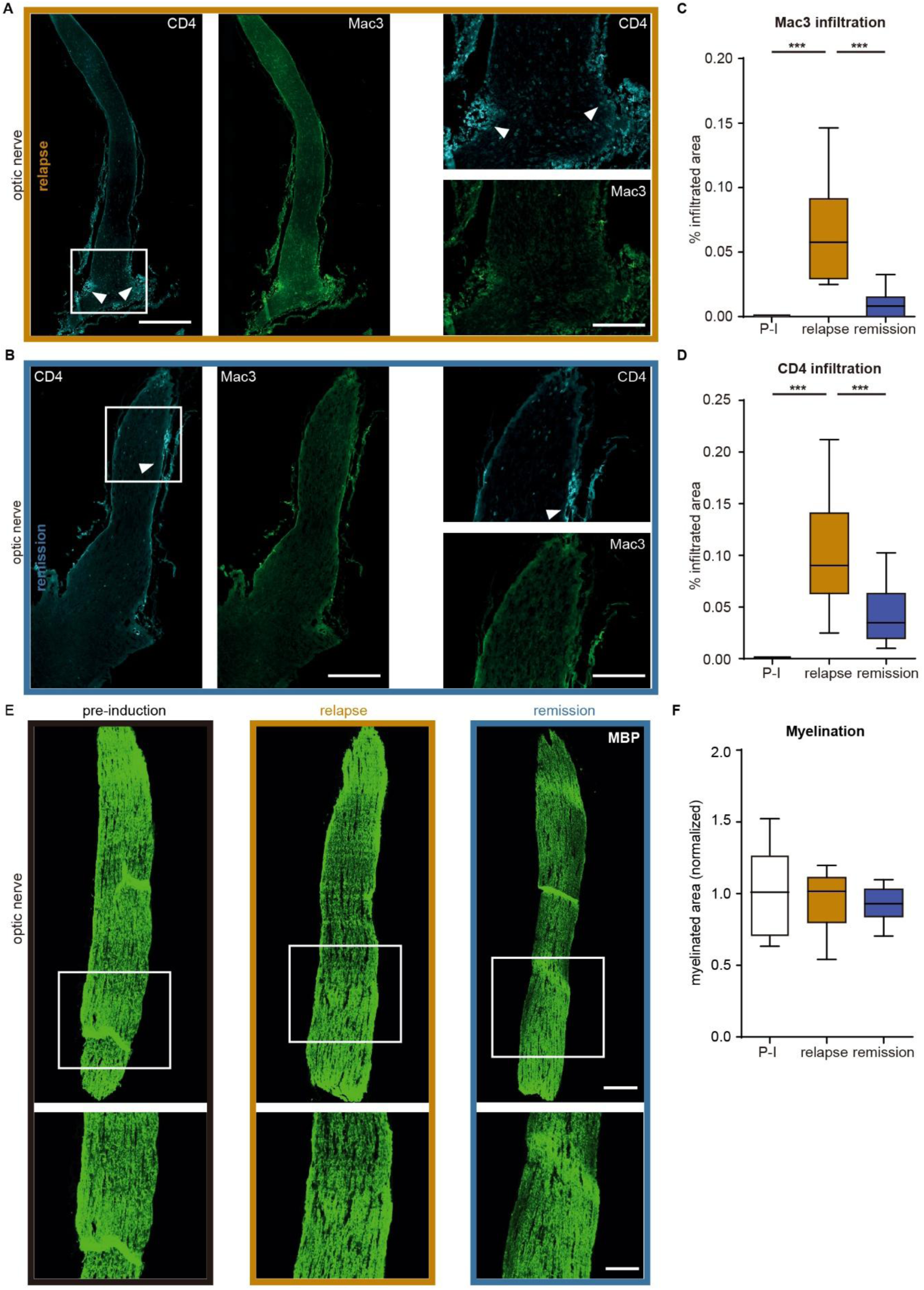
No significant inflammation or demyelination of optic nerve in EAE remission. **A, B:** Immunostaining for CD4 T cells and activated macrophages (Mac3) in optic nerve during **A:** EAE relapse and **B:** remission. **C, D:** Statistics show in relapse a significant infiltration of inflammatory Mac3 cells (**C**, mean ± SD, control 0, relapse 0.06 ± 0.04, remission 0.01 ± 0.01, \*\*\*\**p* < 0.0001) and CD4^+^ T cells (**D**, mean ± SD, CD4 infiltrated area, control 0, relapse 0.10 ± 0.06, remission 0.04 ± 0.03, \*\*\**p* = 0.0003). **E:** MBP myelin staining of the optic nerve. **F:** In relapse and remission we observed no significant demyelination: myelinated area, control 1.0 ± 0.33, relapse 0.95 ± 0.23, remission 0.93 ± 0.13, *p* = 0.72. P values were determined using one-way ANOVA test followed by Bonferonis post-hoc test (\**p* < 0.05, \*\*\**p* < 0.001). Scale bars: overview images 500 µm and inserts 100 µm. *n* = 4-5 animals and 2-4 slices per group.

**Supplementary Figure 6.**
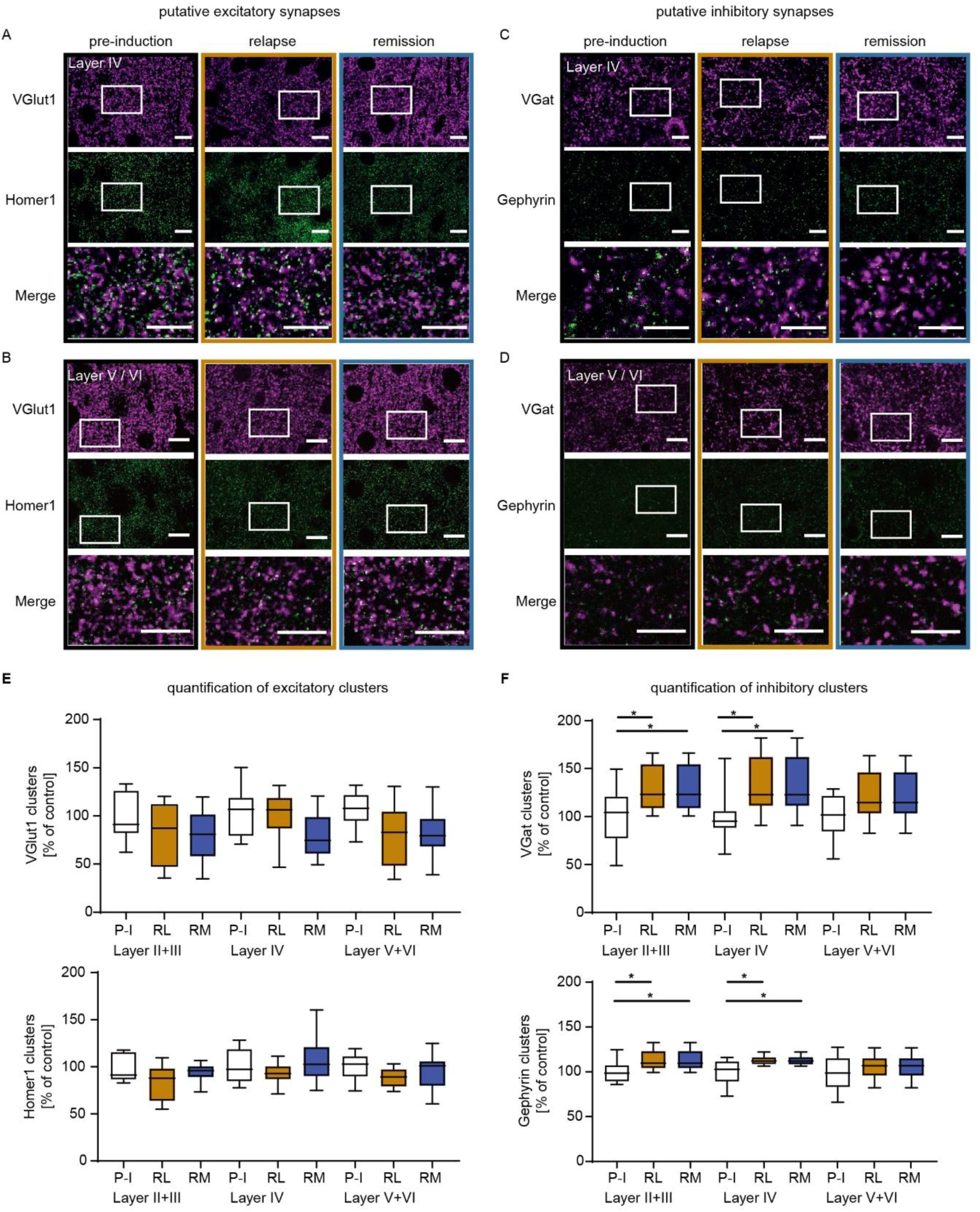
Number of excitatory and inhibitory synapses in Layer II + III, IV and V + VI. **A**. Example pictures of pre- and postsynaptic clusters in pre-induction (P-I), relapse (RL) and remission (RM) mice. Shown is Layer IV. The 3rd row shows merged images which represent excitatory synapses (selections are indicated in the pictures above). **B.** Example pictures of pre- and postsynaptic clusters in P-I, RL and RM mice. Shown is Layer V + VI. The 3rd row shows merged images which represent excitatory synapses (selections are indicated in the pictures above). **C.** Example pictures of pre- and postsynaptic clusters in P-I, RL and RM mice. Shown is Layer IV. The 3rd column shows merged images which represent inhibitory synapses (selections are indicated in the pictures above). **D.** Example pictures of pre-and postsynaptic clusters in P-I, RL and RM mice. Shown is Layer V + VI. The 3rd row shows merged images which represent inhibitory synapses (Selections are indicated in the pictures above). **E.** Single analysis of VGlut1 and Homer1 clusters in Layer II + III, Layer IV and Layer V + VI. **F.** Single analysis of VGat and Gephyrin clusters in Layer II + III, Layer IV and Layer V + VI. Quantification shows that single VGat and Gephyrin clusters are increased in Layer II + III and Layer IV in both RL and RM mice.

**Supplementary Figure 7.**
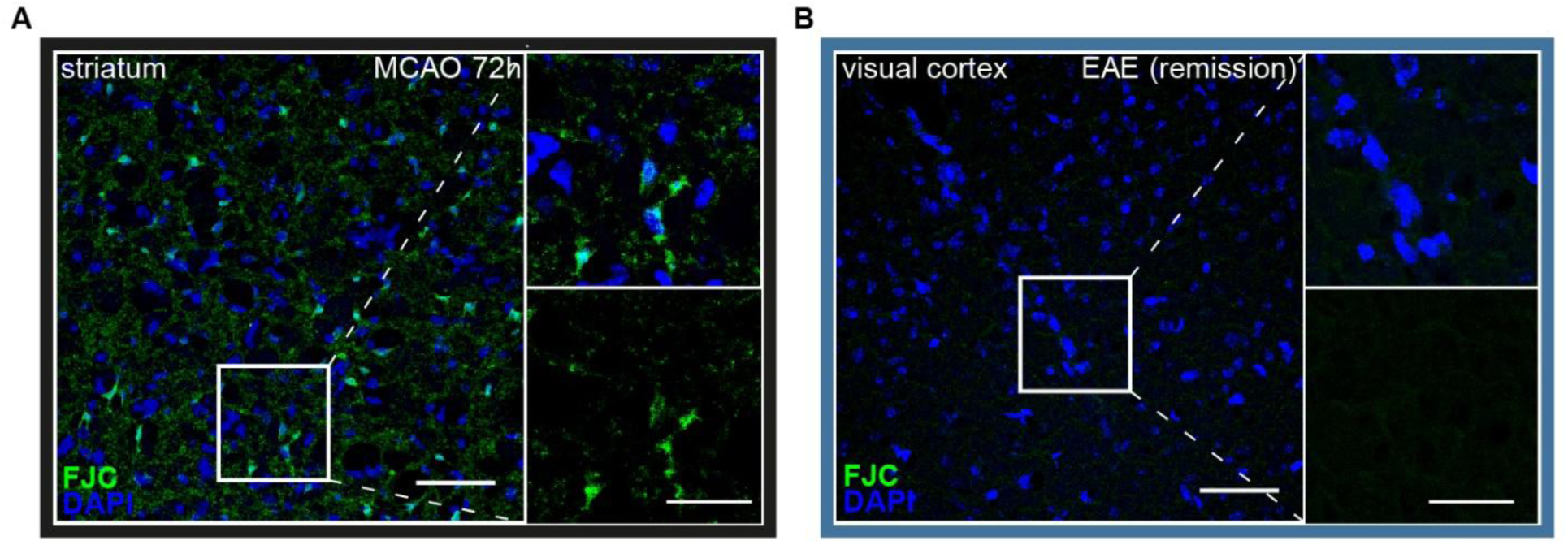
Neuronal cell death staining reveals no widespread neurodegeneration in EAE. **A:** Neurodegeneration indicated by Fluoro Jade C (FJC) positive cells (green) in the ipsilateral hemisphere 72h after Middle Cerebral Artery Occlusion (MCAO). **B:** No Fluoro Jade C-positive cells were detected in EAE in remission. Scale bars 50 µm, overview image (left) was taken at 40x magnification, zoomed-in picture (right) at 63x.

**Supplementary Figure 8:**
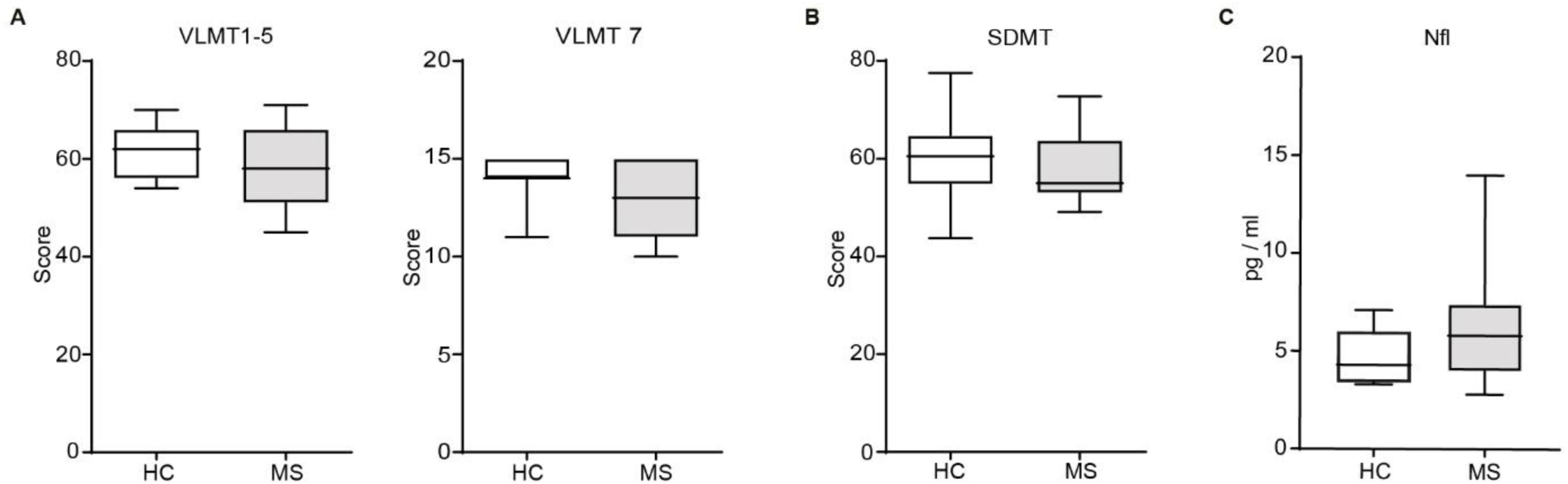
Test battery for patients and controls beside visual discrimination and BORP. **A**: Verbal learning (VLMT 1-5) and long-term memory (VLMT 7, mean ± SD, RRMS: 12.6 ± 2.2; controls: 13.8 ± 1.5, p = 0.059, Mann-Whitney Test) were equal between both groups. **B:** no difference in concentration (SDMT) between MS patients and healthy controls. **C:** NfL values were slightly but not significantly increased in RRMS (in pg / ml, mean ± SD, RRMS: 9.2 ± 13.1; controls: 4.8 ± 1.6, p = 0.08, Mann-Whitney Test).

## References

Aedo-Jury, F., Schwalm, M., Hamzehpour, L. & Stroh, A. 2020. Brain states govern the spatio-temporal dynamics of resting-state functional connectivity. eLife, 9, e53186.

Arnoux, I., Willam, M., Griesche, N., Krummeich, J., Watari, H., Offermann, N., Weber, S., Narayan Dey, P., Chen, C., Monteiro, O., Buettner, S., Meyer, K., Bano, D., Radyushkin, K., Langston, R., Lambert, J. J., Wanker, E., Methner, A., Krauss, S., Schweiger, S. & Stroh, A. 2018. Metformin reverses early cortical network dysfunction and behavior changes in Huntington’s disease. eLife, 7, e38744.

Backhaus, H., Ruffini, N., Wierczeiko, A. & Stroh, A. 2022. An all-optical physiology pipeline towards highly specific and artefact-free circuit mapping *In:* Stroh, A. (ed.) Neuromethods: All-Optical Methods to study Neuronal Activity. Springer.

Bakker, A., Krauss, G. L., Albert, M. S., Speck, C. L., Jones, L. R., Stark, C. E., Yassa, M. A., Bassett, S. S., Shelton, A. L. & Gallagher, M. 2012. Reduction of hippocampal hyperactivity improves cognition in amnestic mild cognitive impairment. Neuron, 74, 467–74.

Benedict, R. H., Deluca, J., Phillips, G., Larocca, N., Hudson, L. D., Rudick, R. & Consortium, M. S. O. A. 2017. Validity of the Symbol Digit Modalities Test as a cognition performance outcome measure for multiple sclerosis. Multiple Sclerosis Journal, 23, 721–733.

Bierbrauer, A., Kunz, L., Gomes, C. A., Luhmann, M., Deuker, L., Getzmann, S., Wascher, E., Gajewski, P. D., Hengstler, J. G., Fernandez-Alvarez, M., Atienza, M., Cammisuli, D. M., Bonatti, F., Pruneti, C., Percesepe, A., Bellaali, Y., Hanseeuw, B., Strange, B. A., Cantero, J. L. & Axmacher, N. 2020. Unmasking selective path integration deficits in Alzheimer&#x2019;s disease risk carriers. Science Advances, 6, eaba1394.

Blumenstock, S. & Dudanova, I. 2020. Cortical and Striatal Circuits in Huntington’s Disease. Front Neurosci, 14, 82.

Boes, A. D., Prasad, S., Liu, H., Liu, Q., Pascual-Leone, A., Caviness, V. S., Jr. & Fox, M. D. 2015. Network localization of neurological symptoms from focal brain lesions. Brain, 138, 3061–75.

Burgold, J., Schulz-Trieglaff, E. K., Voelkl, K., Gutiérrez-Ángel, S., Bader, J. M., Hosp, F., Mann, M., Arzberger, T., Klein, R., Liebscher, S. & Dudanova, I. 2019. Cortical circuit alterations precede motor impairments in Huntington’s disease mice. Scientific Reports, 9, 6634.

Busche, M. A., Chen, X., Henning, H. A., Reichwald, J., Staufenbiel, M., Sakmann, B. & Konnerth, A. 2012. Critical role of soluble amyloid-&#x3b2; for early hippocampal hyperactivity in a mouse model of Alzheimer&#x2019;s disease. Proceedings of the National Academy of Sciences, 109, 8740–8745.

Caputi, N., Matrella, A., Totaro, R., Raparelli, C., Pontecorvo, S., Di Giacomo, D. & Passafiume, D. 2017. Object decision and multiple sclerosis: a preliminary study. Funct Neurol, 32, 69–75.

Charalambous, T., Clayden, J. D., Powell, E., Prados, F., Tur, C., Kanber, B., Chard, D., Ourselin, S., Wheeler-Kingshott, C., Thompson, A. J. & Toosy, A. T. 2020. Disrupted principal network organisation in multiple sclerosis relates to disability. Sci Rep, 10, 3620.

Di Filippo, M., Portaccio, E., Mancini, A. & Calabresi, P. 2018. Multiple sclerosis and cognition: synaptic failure and network dysfunction. Nat Rev Neurosci, 19, 599–609.

Disanto, G., Zecca, C., Maclachlan, S., Sacco, R., Handunnetthi, L., Meier, U. C., Simpson, A., Mcdonald, L., Rossi, A., Benkert, P., Kuhle, J., Ramagopalan, S. V. & Gobbi, C. 2018. Prodromal symptoms of multiple sclerosis in primary care. Ann Neurol, 83, 1162–1173.

Ellwardt, E., Muthuraman, M., Gonzalez-Escamilla, G., Chirumamilla, V. C., Luessi, F., Bittner, S., Zipp, F., Groppa, S. & Fleischer, V. 2022. Network alterations underlying anxiety symptoms in early multiple sclerosis. J Neuroinflammation, 19, 119.

Ellwardt, E., Pramanik, G., Luchtman, D., Novkovic, T., Jubal, E. R., Vogt, J., Arnoux, I., Vogelaar, C. F., Mandal, S., Schmalz, M., Barger, Z., Ruiz De Azua, I., Kuhlmann, T., Lutz, B., Mittmann, T., Bittner, S., Zipp, F. & Stroh, A. 2018. Maladaptive cortical hyperactivity upon recovery from experimental autoimmune encephalomyelitis. Nat Neurosci, 21, 1392–1403.

Engel, S. & Zipp, F. 2022. Preventing disease progression in multiple sclerosis—insights from large real-world cohorts. Genome Medicine, 14, 41.

Farnsworth, D. 1957. The Farnsworth-Munsell 100 Hue Test Manual (revised ed). *Baltimore*: Munsell Color Company.

Fox, M. D. 2018. Mapping Symptoms to Brain Networks with the Human Connectome. N Engl J Med, 379, 2237–2245.

Fu, T., Arnoux, I., Doring, J., Backhaus, H., Watari, H., Stasevicius, I., Fan, W. & Stroh, A. 2021. Exploring two-photon optogenetics beyond 1100 nm for specific and effective all-optical physiology. iScience, 24, 102184.

Gilbert, C. D. & Wiesel, T. N. 1983. Functional Organization of the Visual Cortex. *In:* Changeux, J. P., Glowinski, J., Imbert, M. & Bloom, F. E. (eds.) Progress in Brain Research. Elsevier.

Govier-Cole, A. E., Wood, R. J., Fletcher, J. L., Gonsalvez, D. G., Merlo, D., Cate, H. S., Murray, S. S. & Xiao, J. 2019. Inhibiting Bone Morphogenetic Protein 4 Type I Receptor Signaling Promotes Remyelination by Potentiating Oligodendrocyte Differentiation. eNeuro, 6.

Grienberger, C. & Konnerth, A. 2012. Imaging calcium in neurons. Neuron, 73, 862–85.

Guillery, R. W. & Sherman, S. M. 2002. Thalamic relay functions and their role in corticocortical communication: generalizations from the visual system. Neuron, 33, 163–75.

Guimarães Backhaus, R., Fu, T., Backhaus, H. & Stroh, A. 2021. Pipeline for 2-photon all-optical physiology in mouse: From viral titration and optical window implantation to binarization of calcium transients. STAR Protocols, 2, 101010.

Helmstaedter, C. & Durwen, H. 1990. VLMT: Verbaler Lern-und Merkfähigkeitstest: Ein praktikables und differenziertes Instrumentarium zur Prüfung der verbalen Gedächtnisleistungen. *Schweizer* Archiv für Neurologie, Neurochirurgie und Psychiatrie.

Hubel, D. H. & Wiesel, T. N. 1962. Receptive fields, binocular interaction and functional architecture in the cat’s visual cortex. The Journal of Physiology, 160, 106–154.

Ibbotson, M. & Jung, Y. J. 2020. Origins of Functional Organization in the Visual Cortex. Frontiers in Systems Neuroscience, 14.

Jafari, M., Schumacher, A.-M., Snaidero, N., Ullrich Gavilanes, E. M., Neziraj, T., Kocsis-Jutka, V., Engels, D., Jürgens, T., Wagner, I., Weidinger, J. D. F., Schmidt, S. S., Beltrán, E., Hagan, N., Woodworth, L., Ofengeim, D., Gans, J., Wolf, F., Kreutzfeldt, M., Portugues, R., Merkler, D., Misgeld, T. & Kerschensteiner, M. 2021a. Phagocyte-mediated synapse removal in cortical neuroinflammation is promoted by local calcium accumulation. Nature Neuroscience, 24, 355–367.

Jafari, M., Schumacher, A. M., Snaidero, N., Ullrich Gavilanes, E. M., Neziraj, T., Kocsis-Jutka, V., Engels, D., Jurgens, T., Wagner, I., Weidinger, J. D. F., Schmidt, S. S., Beltran, E., Hagan, N., Woodworth, L., Ofengeim, D., Gans, J., Wolf, F., Kreutzfeldt, M., Portugues, R., Merkler, D., Misgeld, T. & Kerschensteiner, M. 2021b. Phagocyte-mediated synapse removal in cortical neuroinflammation is promoted by local calcium accumulation. Nat Neurosci, 24, 355–367.

Jandric, D., Lipp, I., Paling, D., Rog, D., Castellazzi, G., Haroon, H., Parkes, L., Parker, G. J. M., Tomassini, V. & Muhlert, N. 2021. Mechanisms of Network Changes in Cognitive Impairment in Multiple Sclerosis. Neurology, 97, e1886–e1897.

Kappos, L., Wolinsky, J. S., Giovannoni, G., Arnold, D. L., Wang, Q., Bernasconi, C., Model, F., Koendgen, H., Manfrini, M., Belachew, S. & Hauser, S. L. 2020. Contribution of Relapse-Independent Progression vs Relapse-Associated Worsening to Overall Confirmed Disability Accumulation in Typical Relapsing Multiple Sclerosis in a Pooled Analysis of 2 Randomized Clinical Trials. JAMA Neurol, 77, 1132–1140.

Krishnamoorthy, G. & Wekerle, H. 2009. EAE: an immunologist’s magic eye. Eur J Immunol, 39, 2031–5.

Laatu, S., Revonsuo, A., Hamalainen, P., Ojanen, V. & Ruutiainen, J. 2001. Visual object recognition in multiple sclerosis. J Neurol Sci, 185, 77–88.

Lublin, F. D., Haring, D. A., Ganjgahi, H., Ocampo, A., Hatami, F., Cuklina, J., Aarden, P., Dahlke, F., Arnold, D. L., Wiendl, H., Chitnis, T., Nichols, T. E., Kieseier, B. C. & Bermel, R. A. 2022. How patients with multiple sclerosis acquire disability. Brain.

Luttjohann, A., Rauber, S., Korsen, M., Willison, A. G., Elben, S., Schroeter, C. B., Ruck, T., Albrecht, P., Pawlitzki, M., Stroh, A., Melzer, N., Budde, T. & Meuth, S. G. 2026. Altered network recruitment in multiple sclerosis patients during resting state. Mult Scler Relat Disord, 109, 107111.

Melloni, L., Molina, C., Pena, M., Torres, D., Singer, W. & Rodriguez, E. 2007. Synchronization of Neural Activity across Cortical Areas Correlates with Conscious Perception. The Journal of Neuroscience, 27, 2858–2865.

Menascu, S., Stern, M., Aloni, R., Kalron, A., Magalshvili, D. & Achiron, A. 2019. Assessing cognitive performance in radiologically isolated syndrome. Mult Scler Relat Disord, 32, 70–73.

Murphy, R., O’donoghue, S., Counihan, T., Mcdonald, C., Calabresi, P. A., Ahmed, M. A., Kaplin, A. & Hallahan, B. 2017. Neuropsychiatric syndromes of multiple sclerosis. J Neurol Neurosurg Psychiatry, 88, 697–708.

Parmenter, B., Weinstock-Guttman, B., Garg, N., Munschauer, F. & Benedict, R. H. 2007. Screening for cognitive impairment in multiple sclerosis using the Symbol Digit Modalities Test. Multiple Sclerosis Journal, 13, 52–57.

Polman, C. H., Reingold, S. C., Banwell, B., Clanet, M., Cohen, J. A., Filippi, M., Fujihara, K., Havrdova, E., Hutchinson, M., Kappos, L., Lublin, F. D., Montalban, X., O’connor, P., Sandberg-Wollheim, M., Thompson, A. J., Waubant, E., Weinshenker, B. & Wolinsky, J. S. 2011. Diagnostic criteria for multiple sclerosis: 2010 revisions to the McDonald criteria. Ann Neurol, 69, 292–302.

Putcha, D., Brickhouse, M., O’keefe, K., Sullivan, C., Rentz, D., Marshall, G., Dickerson, B. & Sperling, R. 2011. Hippocampal Hyperactivation Associated with Cortical Thinning in Alzheimer’s Disease Signature Regions in Non-Demented Elderly Adults. The Journal of Neuroscience, 31, 17680–17688.

Riddoch, M. J. & Humphreys, G. W. 1993. Birmingham object recognition battery, Lawrence Erlbaum Associates.

Rochefort, Nathalie l., Narushima, M., Grienberger, C., Marandi, N., Hill, Daniel n. & Konnerth, A. 2011. Development of Direction Selectivity in Mouse Cortical Neurons. Neuron, 71, 425–432.

Rosales Jubal, E., Schwalm, M., Dos Santos Guilherme, M., Schuck, F., Reinhardt, S., Tose, A., Barger, Z., Roesler, M. K., Ruffini, N., Wierczeiko, A., Schmeisser, M. J., Schmitt, U., Endres, K. & Stroh, A. 2021. Acitretin reverses early functional network degradation in a mouse model of familial Alzheimer’s disease. Scientific Reports, 11, 6649.

Rosenthal, Z. P., Raut, R. V., Yan, P., Koko, D., Kraft, A. W., Czerniewski, L., Acland, B., Mitra, A., Snyder, L. H., Bauer, A. Q., Snyder, A. Z., Culver, J. P., Raichle, M. E. & Lee, J. M. 2020. Local Perturbations of Cortical Excitability Propagate Differentially Through Large-Scale Functional Networks. Cereb Cortex, 30, 3352–3369.

Siffrin, V., Radbruch, H., Glumm, R., Niesner, R., Paterka, M., Herz, J., Leuenberger, T., Lehmann, S. M., Luenstedt, S., Rinnenthal, J. L., Laube, G., Luche, H., Lehnardt, S., Fehling, H. J., Griesbeck, O. & Zipp, F. 2010. In vivo imaging of partially reversible th17 cell-induced neuronal dysfunction in the course of encephalomyelitis. Immunity, 33, 424–36.

Singer, W. 2013. Cortical dynamics revisited. Trends in Cognitive Sciences, 17, 616–626.

Sjogard, M., Wens, V., Van Schependom, J., Costers, L., D’hooghe, M., D’haeseleer, M., Woolrich, M., Goldman, S., Nagels, G. & De Tiege, X. 2021. Brain dysconnectivity relates to disability and cognitive impairment in multiple sclerosis. Hum Brain Mapp, 42, 626–643.

Stroh, A., Schweiger, S., Ramirez, J. M. & Tuscher, O. 2024. The selfish network: how the brain preserves behavioral function through shifts in neuronal network state. Trends Neurosci, 47, 246–258.

Tchumatchenko, T., Geisel, T., Volgushev, M. & Wolf, F. 2010. Signatures of synchrony in pairwise count correlations. Frontiers in Computational Neuroscience, 4.

Tetreault, A. M., Phan, T., Orlando, D., Lyu, I., Kang, H., Landman, B., Darby, R. R. & Alzheimer’s Disease Neuroimaging, I. 2020. Network localization of clinical, cognitive, and neuropsychiatric symptoms in Alzheimer’s disease. Brain, 143, 1249–1260.

Tur, C., Carbonell-Mirabent, P., Cobo-Calvo, A., Otero-Romero, S., Arrambide, G., Midaglia, L., Castillo, J., Vidal-Jordana, A., Rodriguez-Acevedo, B., Zabalza, A., Galan, I., Nos, C., Salerno, A., Auger, C., Pareto, D., Comabella, M., Rio, J., Sastre-Garriga, J., Rovira, A., Tintore, M. & Montalban, X. 2023. Association of Early Progression Independent of Relapse Activity With Long-term Disability After a First Demyelinating Event in Multiple Sclerosis. JAMA Neurol, 80, 151–160.

Turrigiano, G. 2012. Homeostatic synaptic plasticity: local and global mechanisms for stabilizing neuronal function. Cold Spring Harb Perspect Biol, 4, a005736.

Urai, A. E., Aguillon-Rodriguez, V., Laranjeira, I. C., Cazettes, F., International Brain, L., Mainen, Z. F. & Churchland, A. K. 2021. Citric Acid Water as an Alternative to Water Restriction for High-Yield Mouse Behavior. eNeuro, 8.

Vleugels, L., Lafosse, C., Van Nunen, A., Nachtergaele, S., Ketelaer, P., Charlier, M. & Vandenbussche, E. 2000. Visuoperceptual impairment in multiple sclerosis patients diagnosed with neuropsychological tasks. Mult Scler, 6, 241–54.

Wiendl, H., Gold, R., Berger, T., Derfuss, T., Linker, R., Maurer, M., Aktas, O., Baum, K., Berghoff, M., Bittner, S., Chan, A., Czaplinski, A., Deisenhammer, F., Di Pauli, F., Du Pasquier, R., Enzinger, C., Fertl, E., Gass, A., Gehring, K., Gobbi, C., Goebels, N., Guger, M., Haghikia, A., Hartung, H. P., Heidenreich, F., Hoffmann, O., Kallmann, B., Kleinschnitz, C., Klotz, L., Leussink, V. I., Leutmezer, F., Limmroth, V., Lunemann, J. D., Lutterotti, A., Meuth, S. G., Meyding-Lamade, U., Platten, M., Rieckmann, P., Schmidt, S., Tumani, H., Weber, F., Weber, M. S., Zettl, U. K., Ziemssen, T., Zipp, F. & Multiple Sclerosis Therapy Consensus, G. 2021. Multiple Sclerosis Therapy Consensus Group (MSTCG): position statement on disease-modifying therapies for multiple sclerosis (white paper). Ther Adv Neurol Disord, 14, 17562864211039648.

Yang, Y., Wang, M., Xu, L., Zhong, M., Wang, Y., Luan, M., Li, X. & Zheng, X. 2022. Cerebellar and/or Brainstem Lesions Indicate Poor Prognosis in Multiple Sclerosis: A Systematic Review. Front Neurol, 13, 874388.

Zigmond, A. S. & Snaith, R. P. 1983. The hospital anxiety and depression scale. Acta psychiatrica scandinavica, 67, 361–370.

Zott, B., Busche, M. A., Sperling, R. A. & Konnerth, A. 2018. What Happens with the Circuit in Alzheimer’s Disease in Mice and Humans? Annu Rev Neurosci, 41, 277–297.

